# Timing and magnitude of the type-I interferon response are determinants of disease tolerance in arbovirus infection

**DOI:** 10.1101/2022.10.19.512818

**Authors:** Alexandra Hardy, Siddharth Bakshi, Wilhelm Furnon, Oscar MacLean, Quan Gu, Margus Varjak, Mariana Varela, Muhamad A Aziz, Andrew Shaw, Rute Maria Pinto, Natalia Cameron Ruiz, Catrina Mullan, Aislynn Taggart, Ana Filipe, Richard E. Randall, Sam Wilson, Meredith E. Stewart, Massimo Palmarini

**Author notes:** contributed equally to this study (Siddharth Bakshi, Wilhelm Furnon). Institute of Biological Sciences, Faculty of Science, Universiti Malaya, 50603 Kuala Lumpur, Malaysia (Muhamad A Aziz).

## Abstract

Infected hosts possess two alternative strategies to protect themselves against the negative impact of virus infections: (i) “resistance”, directed to abrogate virus replication, or (ii) “disease tolerance”, aimed to avoid organ and tissue damage without overly controlling viral burden. The overall principles governing pathogen resistance are well understood, while less is known about those involved in disease tolerance. Here, we studied bluetongue virus (BTV), the cause of a major disease of ruminants, bluetongue, as a model system to investigate the mechanisms of disease tolerance. BTV induces clinical disease mainly in sheep, while cattle are considered reservoirs of infection, rarely exhibiting clinical symptoms despite sustained viremia. Here, we show that BTV consistently reaches higher titres in ovine primary cells, compared to cells derived from cattle. The variable replication kinetics of BTV in sheep and cattle cells were mostly abolished by abrogating the cell type-I interferon (IFN) response. By screening a library of bovine interferon stimulated genes (ISGs), we identified restriction factors blocking BTV replication, however both sheep and cattle orthologues of these antiviral genes possess anti-BTV properties. Importantly, we demonstrate that BTV induces a faster host cell protein synthesis shutoff in primary sheep cells, compared to cattle cells, which results in an earlier downregulation of antiviral proteins. Moreover, by RNAseq we also show a more pronounced expression of ISGs in BTV infected cattle cells compared to sheep cells. Our data provide a new perspective on how the type-I IFN response in reservoir species can have overall positive effects on both virus and host evolution.

## INTRODUCTION

Many pathogens infect more than one animal species resulting in dramatically different clinical outcomes as a result of complex differences in pathogen-host interactions [1-3]. The infected host can protect itself from diseases using essentially two distinct strategies: resistance and tolerance [4-7]. Resistance refers to the host immune response’s ability to hamper and eliminate the pathogen once infection has occurred. Conversely, tolerance (also referred to as “resilience” and not to be confused with “immunological tolerance”), is a host defence strategy that protects against the deleterious effects of pathogen infection and the host immune response in the face of high pathogen burden [8-11]. Disease tolerance was first described in the context of the selective pressures provided by parasites and herbivores on plant evolution. Only relatively recently this concept has been explored in the context of pathogen-mammalian host interactions [10, 12, 13]. A variety of factors including genetic traits, innate and adaptive immunity, microbiome and age have been shown to play a role in disease tolerance [14].

The distinction between resistance and tolerance is of paramount importance in understanding the ecology and epidemiology of infectious diseases. Natural “reservoir” species of given pathogens can use disease tolerance as an effective mechanism to cope with infection. At the same time, high pathogen burden and extended infection times can favour onward transmission to target susceptible species. For example, some viral infections (Nipah, Hendra) in different species of bats maintain high viral burden without apparent disease, while infection of these viruses results in severe disease in humans [15]. African buffaloes are tolerant to foot and mouth disease virus relative to domestic cattle, while wild hogs are in general tolerant to African swine fever virus as opposed to domestic pigs [16, 17].

In this study, we aimed to dissect virus-host interactions in susceptible and disease tolerant host species using a tractable experimental system. We used bluetongue virus (BTV), an arbovirus of ruminants with a significant impact on the agriculture sector [18-21]. BTV is a dsRNA virus within the *Reoviridae* family existing in nature as more than 27 serotypes [22-24]. BTV can essentailly infect all domestic and wild ruminant species, but the clinical manifestations of infection can vary considerably, both between species and individuals [25, 26]. BTV infects and induces viraemia both in cattle and sheep, although in general only the latter display the severe clinical signs associated with bluetongue. As such, sheep are recognised as susceptible hosts whilst cattle are mostly asymptomatic [27-31]. Clinical manifestations particularly evident in sheep include fever, nasal discharge, diffuse oedema affecting the head and lungs, as well as haemorrhagic lesions [32-35]. In the infected organs, endothelial cells (EC) have been described as major cellular targets of BTV, where virus replication may cause cell injury [36-39]. Damage to the EC is believed to be crucial in the pathogenesis of the disease. Indeed, injuries of the EC lining the small blood vessels clearly reflect the clinical signs observed during BTV infection. The increased vascular permeability associated with EC damage may account for the observed oedema as well as the reported lesions such as vascular thrombosis and haemorrhages [36, 37, 40]

Disease in cattle is quite rare with only intermittent reports of field cases of bluetongue disease in cattle [28, 41]. In addition, clinical disease has been very difficult to reproduce experimentally in cattle, regardless of animal age, virus serotype or route of inoculation [33, 42-47]. A possible exception is the North European BTV-8 from 2006, which induced one of the largest outbreaks of bluetongue in history with elevated mortality in sheep but also some morbidity in cattle (although of reduced severity compared to sheep)[42].

In this study, we aimed to investigate whether susceptibility or tolerance to BTV could correlate with fundamental difference in virus-host cell interactions that can be assessed in *in vitro* experiments. We focused our studies on primary cells derived from sheep and cattle, the two animal species exhibiting a different clinical outcome to BTV infection. Our findings provide a unique comparative approach to understand how virus-host interactions may influence disease tolerance and susceptibility to virus infections.

## RESULTS

### Replication kinetics of BTV in bovine and ovine primary cells

We first investigated whether host resilience or susceptibility to BTV infection could be correlated to the virus replication kinetics in primary cells collected from sheep (susceptible species) or cattle (resilient species). We carried out virus replication assays in primary skin fibroblasts and endothelial cells. We chose primary cells as they are likely to possess an intact cell autonomous innate immune response unlike many established immortalised cell lines. BTV is an arbovirus gaining access into the infected animals through the skin and skin fibroblasts were a convenient system to use although we have no evidence that they are infected *in vivo*. Endothelial cells are instead known targets of BTV infection *in vivo* [36-39].

We were particularly careful in performing these assays in distinct cell preparations collected from multiple donors to compensate for individual and batch-to-batch variation. In addition, we used two distinct BTV serotypes; BTV-8 and BTV-2, to ensure that differences observed were not associated with a single virus strain. In replication assays, both BTV-8 and BTV-2 reached statistically significant higher titres in OvFib and OvEC compared to the titres reached in cattle cells (BovFib and BovEC) (Fig 1A). Interestingly, both BTV-2 and BTV-8 reached titres at times 100-fold higher in OvEC as opposed to BovEC, while differences of titres between ovine and bovine in fibroblasts were more modest (about 10-fold). The differences we observed in the virus replication kinetics correlated to differences in the progression of cytopathic effect (CPE) during the time course of infection (Fig S1). The monolayer of OvEC was destroyed almost completely by BTV within 72h post-infection while BovEC showed restricted foci of infection that did not progress as extensively even at 96 hpi.

**Figure 1.**
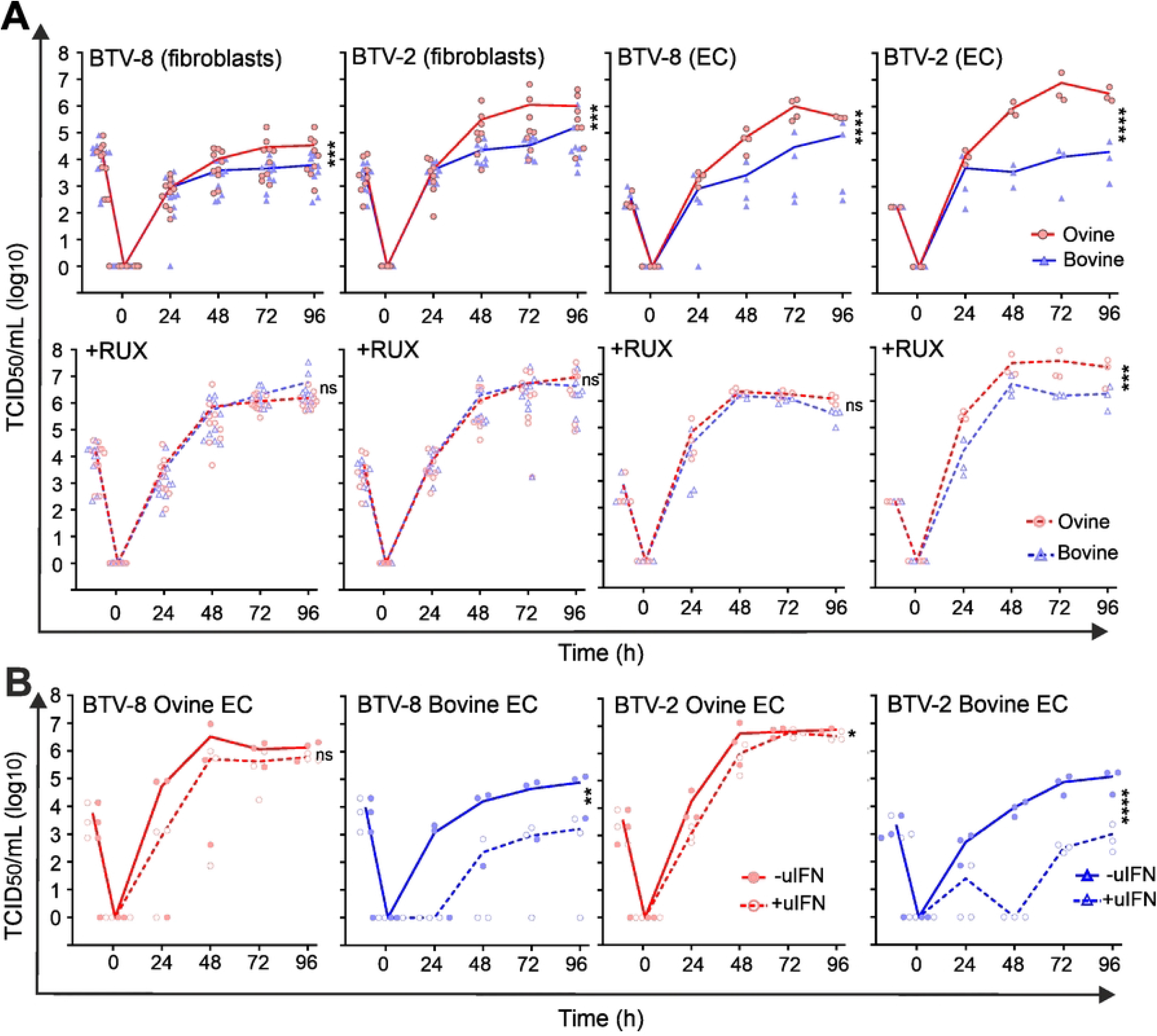
Differences in the replication kinetics of BTV in primary cells is dependent on the type-I IFN response. (**A**) Virus replication curves of BTV-8 and BTV-2 in ovine (red) and bovine (blue) primary fibroblasts and endothelial cells (EC). Datapoints for independent experiments are shown (BTV-8 n=10 and BTV-2 n=8 for ovine and bovine fibroblasts; BTV-8 n=4 and BTV-2 n=3 for ovine and bovine EC). Bottom panels show same replication curves in the presence of 4 μM RUX. Experiments in the presence or absence of RUX were carried out at the same time. (**B**) Virus replication curves of BTV-8 and BTV-2 in ovine (red) and bovine (blue) EC. Cells were mock-treated (solid lines) or pre-treated with 500U of uIFN before infection. Independent experimental repeats for BTV-8 n= 3 and BTV-2 n=3 for ovine and bovine endothelial cells. Both for experiments shown in A and B, cells were infected at MOI of 0.01 and supernatants were harvested at the indicated time post-infection. Cell-free virus was titrated by endpoint dilution on BSR cells. Titres are expressed as TCID_50_/mL and represent an average of independent experiments as indicated using at least three batches of primary cells independently generated. A one-way ANOVA with Holm-Šídák post hoc correction test was carried out on the area under the curve (AUC) values to assess statistical significance between conditions. ns = p>0.05, *= p<0.05, **= p<0.01, *** = p<0.001, **** =p<0.0001.

### Differences in the replication kinetics of BTV in bovine and sheep cells is modulated by the IFN response

BTV is a strong inducer of type I IFN, both *in vivo* and *in vitro* [48-53]. Hence, we investigated whether the antiviral IFN response played a role in the different phenotype displayed by BTV in sheep and cattle cells. To this end, we examined virus replication in the presence of Ruxolitinib (Rux) an efficient inhibitor of the IFN signalling pathway acting on the Janus kinase 1 (JAK1) and JAK2 proteins [54, 55]. Higher BTV titres were observed in cells treated with Rux (Fig 1A, bottom panels), relative to those obtained in the untreated cells at all timepoints and in all cell types used. Strikingly, the differences in the BTV titres after Rux treatment were more substantial in bovine cells. Indeed, the addition of Rux was able to rescue BTV replication in bovine cells to levels comparable to those in Rux treated ovine cells. Hence, the JAK/STAT pathway and the initiation of the signalling cascade leading to the regulation of hundreds of IFN-stimulated genes (ISGs) appears to play a role in the different replication kinetics shown by BTV in ovine and bovine cells.

We next performed virus replication assays in cells with or without pre-treatment with type-I IFN (500 IU/mL uIFN). The induction of an antiviral state restricted BTV-8 replication, with the effect more prominent in cattle as opposed to sheep fibroblasts (Fig 1B). As ECs had showed the largest differences in the BTV replication kinetics between sheep and cattle cells, we concentrated further on examining these differences. The presence of IFN minimally affected BTV-8 or BTV-2 replication in OvEC as titres were ∼15 to 60-fold lower at 24 hpi in cells pre-treated with uIFN compared to untreated cells but there were no major differences at later time-points for both serotypes tested. Conversely, replication of both BTV-8 and BTV-2 was significantly hampered in BovEC pre-treated with uIFN compared to untreated cells. Replication of both BTV-8 and BTV-2 was severely delayed and reached very low titres at the early time points, and only started to increase from 48 hpi onwards. In BovEC, BTV-2 and BTV-8 reached titres ∼40-120 fold higher in mock-treated cells compared to IFN-treated cells (Fig 1B). Hence, the data obtained so far indicate that the nature of the antiviral response may determine whether a host is susceptible or tolerant to BTV induced disease.

### Identification of ISGs restricting BTV replication

The IFN response appeared to suppress BTV replication more effectively in bovine cells. Hence, we next investigated whether ISGs that are induced upon IFN treatment in bovine cells are efficient at blocking BTV replication, while their ovine orthologues may be less effective in doing so. To identify bovine ISGs which could negatively impact BTV replication, we used a medium throughput lentivirus-based overexpression screening platform. This system was previously used to identify antiviral genes acting against a range of viruses [56-59] and relies on lentivirus vectors co-expressing the ISG of interest and a fluorescent protein (RFP). HEK-293 cells were challenged with BTV expressing the green fluorescent protein (GFP), and the level of virus replication in the presence of each individual ISG was assessed via flow cytometry (Fig 2A-B). For our screening purposes, we engineered a reporter BTV expressing a monomeric GFP fused to the NS1 ORF (BTV-8mGFP).

**Figure 2.**
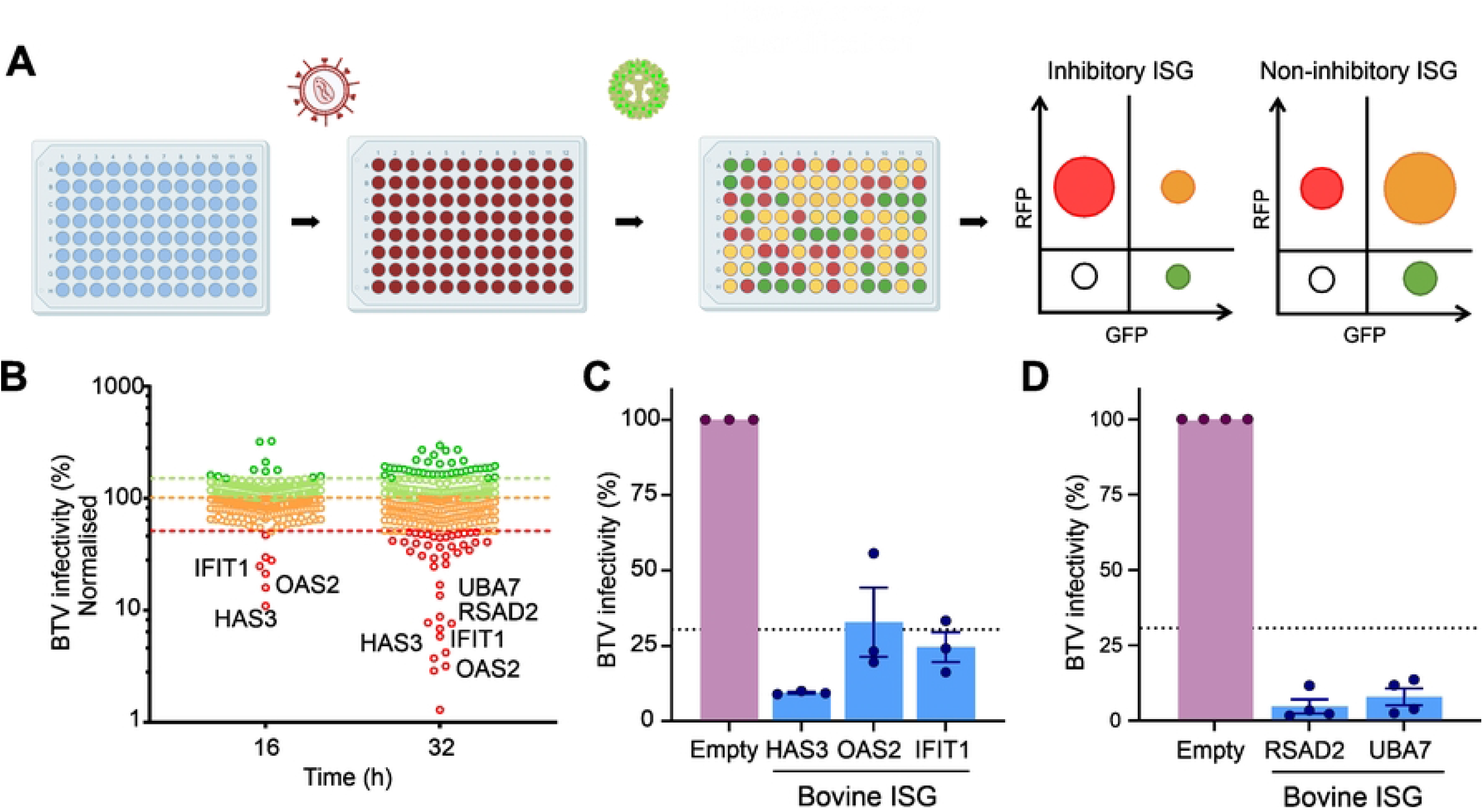
Bovine ISG screen reveals gene candidates with anti-BTV properties. (**A**) A schematic representation of the bovine ISG over-expression screen. 293T cells were transduced with lentiviruses expressing ISGs and a red fluorescent protein (RFP). Cells are subsequently infected with BTV-mGFP. At either 16h or 32 hpi cells were fixed, and number of infected cells determined via flow cytometry. Plots representing the relative importance of different cell populations gated on their expression of GFP (virus-infected cells) or RFP (efficiently transduced cells) for an inhibitory or a non-inhibitory ISG are schematically represented. (**B**) Scatter plot representing the percentage of BTV-mGFP replication for each gene normalised to the median; screen at 16 hpi and 32 hpi. Genes restricting BTV by more than 30% are considered inhibitory. The screen at 32 hpi was undertaken in the presence of 0.625 ug/mL puromycin. (**C**) Validation of antiviral bovine ISG inhibiting BTV replication ay 16 hpi. Data are shown as the mean +/- standard errors of three independent experiments. (**D**) Validation of antiviral bovine ISG inhibiting BTV replication at 32 hpi. Data are shown as the mean +/- standard errors of four independent experiments. Some of the schematic images were obtained from Biorender.com under a laboratory licence.

We designed and constructed a bovine ISG library including “core ISGs” conserved across mammalian evolution and those that are most highly upregulated in response to IFN in bovine primary cells, as identified in one of our previous studies (Supplementary Table 1) [60]. We carried out the screens in HEK-293T cells transduced with the lentivirus library and infected cells were fixed at two different times post-infection (16 and 32 hpi) and then analysed by flow cytometry. By establishing an arbitrary cut-off value of 30% for the normalised level of BTV replication, we identified 17 potential hits with anti-BTV-8mGFP properties: 7 genes in the 16 hpi screen and 15 in the 32 hpi screen (Fig 2B; Supplemental Table1). Six of the 7 genes identified in the 16 hpi screen were also identified as restrictive at 32 hpi. Given our experience with ISG screens against a variety of viruses [56-59], we ruled out from subsequent analyses ISGs with (i) known signalling properties (IRF2, IFIH1, DDX58), or (ii) able to induce expression of a reporter gene under the control of an ISRE (interferon stimulated response element) promoter in the same type of assays (IRF-1, CDADC1, CXCR4, ISG20, STARD8), or (iii) known to affect the general cell metabolism (IDO1) [61]. We also excluded SLFN12L as it was identified only in the 16 hpi timepoint and not confirmed in the 32 hpi screen.

We validated these results using different lentivirus preparations) with the remaining hits: HAS3 (Hyaluronan synthase 3), IFIT1 (Interferon induced protein with tetratricopeptide repeats 1), OAS2 (2’-5’-Oligoadenylate synthetase 2), RSAD2 [Radical S-adenosyl methionine domain containing 2] and UBA7 [Ubiquitin like modifier activating enzyme 7], which confirmed their anti-BTV properties. All these ISGs were confirmed to possess anti-BTV properties in the validation assays (Fig. 2C-D).

### Ablation of IFIT1 and RSAD2 reduces BTV sensitivity to IFN

IFIT1 and RSAD2 are core ISGs conserved across vertebrate evolution and possess broad antiviral activity [60]. Furthermore, sheep RSAD2 was shown in a previous study to possess anti-BTV activity [62]. To complement the data obtained with our overexpression approach, we used CRISPR/Cas9 to knock-out (KO) IFIT1 and RSAD2 expression and determine their effect on BTV replication. While we could not successfully generate KO cells with the CRISPR/Cas9 system using primary cells, we were successful in immortalised primary bovine fibroblast (BovFibT). We confirmed the ablation of IFIT1 and RSAD2 by immunoblotting in two clonally selected KO cell populations after stimulation with uIFN for 24h. The expression of IFIT1 and RSAD2 was either undetectable or greatly reduced in the KO cells in comparison to the parental cells (Fig 3A, B). Importantly, induction of pSTAT1 in response to uIFN stimulation in KO and parental cells displayed no major differences. These data therefore suggest that the ability of the KO cells to respond to IFN was maintained, although we noticed in the RSAD2-KO cells a somewhat reduced pSTAT1 signal, possibly reflecting clonal variability (Fig 3B).

**Figure 3.**
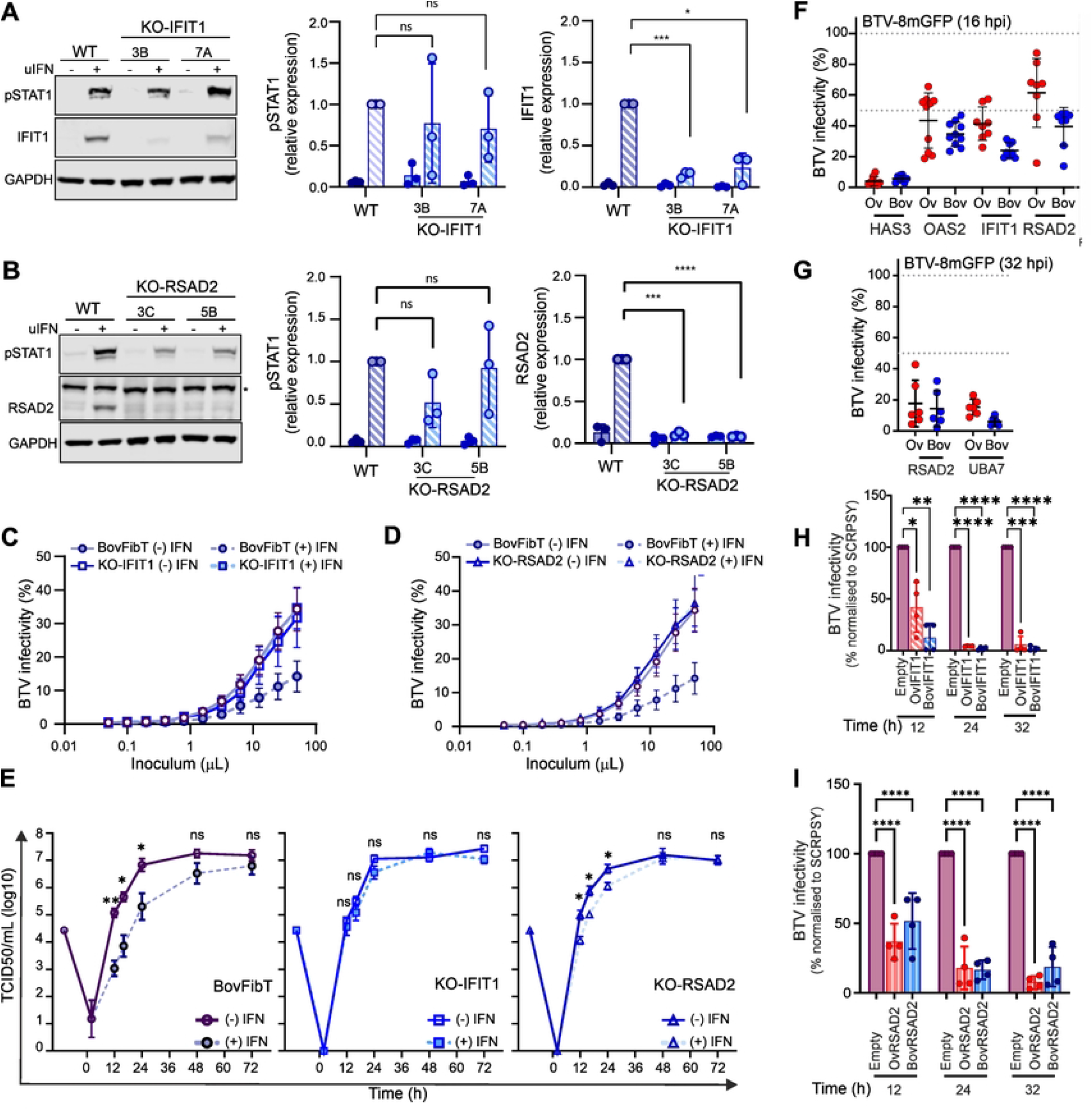
BTV replication is restricted by both cattle antiviral ISGs and their sheep orthologues. (**A**) (Left) Western blotting of KO-IFIT1 cells (clones 3B and 7A) and parental BovFibT cells (WT) showing expression of IFIT1, phosphorylated STAT1 and GAPDH after IFN treatment. Right panels show quantification of western blots using the Image Studio Lite software (LI-COR Biosciences). Data were obtained by from 3 independent experiments. (**B**) (Left) Western blotting of KO-RSAD2 cells (clones 3C and 5B) and parental BovFibT cells (WT) showing expression of RSAD2, phosphorylated STAT1 and GAPDH after IFN treatment. Right panels show quantification of western blots using the Image Studio Lite software (LI-COR Biosciences). Data were obtained by from 3 independent experiments. (**C**) Graph showing infectivity of BTV-8 mGFP in KO-IFIT1 and parental BovFibT in cells pre-treated with IFN (1000 U), or carrier control, before infection with serial dilutions of BTV8-mGFP for 6 h. Cells were then fixed and mGFP expression was determined by flow cytometry. (**D**) Graph showing infectivity of BTV-8 mGFP in KO-RSAD2 and parental BovFibT in cells pre-treated with IFN (1000 U), or carrier control, before infection with serial dilutions of BTV8-mGFP for 6 h. Cells were then fixed and mGFP expression was determined by flow cytometry. (**E**) Virus replication curves of growth of BTV-8 in immortalised BovEC (left), KO-IFIT1 and KO-RSAD2 cells in the presence (dashed lines) or absence (solid lines) of 500U uIFN pre-treatment. Cells were infected at MOI of 0.01 and supernatants were harvested at the indicated time points post-infection. Cell-free virus was titrated by endpoint dilution on BSR cells, and titres are expressed as TCID50/mL and represent mean +/- SEM of at least 3 independent experiments. Statistical significance between –uIFN and +uIFN conditions for each timepoint was calculated using t-tests with Welch’s correction. (**F-G**) Relative infectivity of BTV-8mGFP in 293T cells overexpressing bovine and ovine restriction factors at 16 (F) and 32 (G) hpi (n=8). 293T cells were transduced with lentiviruses expressing the ovine or bovine orthologues for 48 h, infected with BTV-8mGFP for 16 or 32 h, and mGFP expression was quantified by flow cytometry. BTV infectivity was normalised to the mean obtained from all the negative control wells. Each dot represents an independent repeat. The mean and standard deviation are represented for each condition. (**H-I**) Virus titres of BTV-8 in CPT-Tert stably expressing either ovine or bovine IFIT1 (H), and ovine or bovine RSAD2 (I) at the times indicated. Cells stably transfected with an empty lentivirus was used as control. Cells were infected with BTV-8 (MOI ∼ 0.01) and supernatants were harvested at the indicated time points post-infection. Cell-free virus titres were quantified by endpoint dilution and normalised to titres obtained from the control cell line. Data (n =4) is from 2 independent BTV-8 stocks and 2 independently generated stable cell line for each gene tested. Multiple t-tests were carried out following a Shapiro-Wilk normality test.

We next evaluated the effects of ISGs on the initial stages of viral infection using BTV-8mGFP. IFIT1-KO, RSAD2-KO and parental BovFibT were stimulated with uIFN and then infected with serial dilutions of BTV8-mGFP for 6 hours. As expected, we observed a clearly visible decrease in the number of positive mGFP cells (a proxy for BTV infection) in IFN-treated cells, as opposed to mock-treated cells (Fig 3C-D). However, we observed no differences in the number of BTV8-mGFP infected KO cells whether they were pre-treated with IFN or mock-treated (Fig 3C-D).

To further assess the effect of IFIT1 and RSAD2 knock-out on BTV replication, BovFibT, IFIT-KO and RSAD2-KO were either stimulated with uIFN or mock-treated for 24 h prior to infection with wild type BTV-8 (MOI of 0.01) (Fig 3E). As expected, pre-treatment of BovFibT cells with IFN suppressed the replication kinetics of BTV-8. On the other hand, pre-treatment of IFIT1-KO cells did not affect the replication kinetics of BTV-8. Conversely, IFN stimulation of the RSAD2-KO cells resulted in an intermediate phenotype with differences in the BTV-8 replication kinetics that were statistically significant compared to untreated cells, but these differences were not as large as those observed in the parental BovFibT. These results provide evidence that the IFN response is significantly less efficient at controlling BTV replication in the absence of IFIT1 and RSAD2.

### No difference in antiviral activity displayed by sheep and cattle ISG orthologues possessing anti-BTV activity

Our data thus far showed that BTV replication in bovine cells is hampered by the host IFN response. In addition, we have also identified at least five bovine ISGs restricting BTV replication. Hence, we next compared the antiviral activity of the ovine and bovine ISG orthologues. We used the same exogenous expression assays described above, using independently generated lentiviruses encoding for either the ovine or bovine orthologues of IFIT1, RSAD2, OAS2, HAS3 and UBA7. Overall, we did not observe significant differences in the ability of sheep and bovine ISG orthologues to restrict BTV replication at either 16 hpi (Fig 3F) or 32 hpi (Fig 3G).

As the above experiments were undertaken in a human cell line with a reporter virus, we further validated the results obtained using a sheep cell line (CPT-Tert) stably expressing either IFIT1 or RSAD2. As in the previous experiments, both sheep and bovine IFIT1 and RSAD2 significantly restricted BTV replication (Fig 3H-I).

### Relative impairment of BTV replication in BoVEC is detectable as early as 4 hours post-infection

Next, we further dissected the early events of the BTV life cycle using BTV-8mGFP to better assess small differences in the early events of BTV replication in BovEC and OvEC. We synchronised infection of BovEC and OvEC and incubated cells for either 2, 4 or 6 hpi before flow cytometry. At 2 hpi there was no significant difference in the number of infected OvEC and BovEC, but by 4 hpi there was a significant increase in the number of BTV-mGFP infected OvEC cells (Fig 4A). The mean fluorescent intensity of the two cell types was similar at 2 and 4 hpi while it became significantly higher in infected OvEC at 6 hpi, suggesting greater translation of BTV proteins in these cells at this time point (Fig 4B). We repeated the experiments described above also in immortalised ovine and bovine EC (OvEChTert and BovEChTert) and obtained similar results (Fig 4C-D). These data suggest that the differences in the replication kinetics of BTV in OvEC and BovEC can be detected at least as early as 4 hpi.

**Figure 4.**
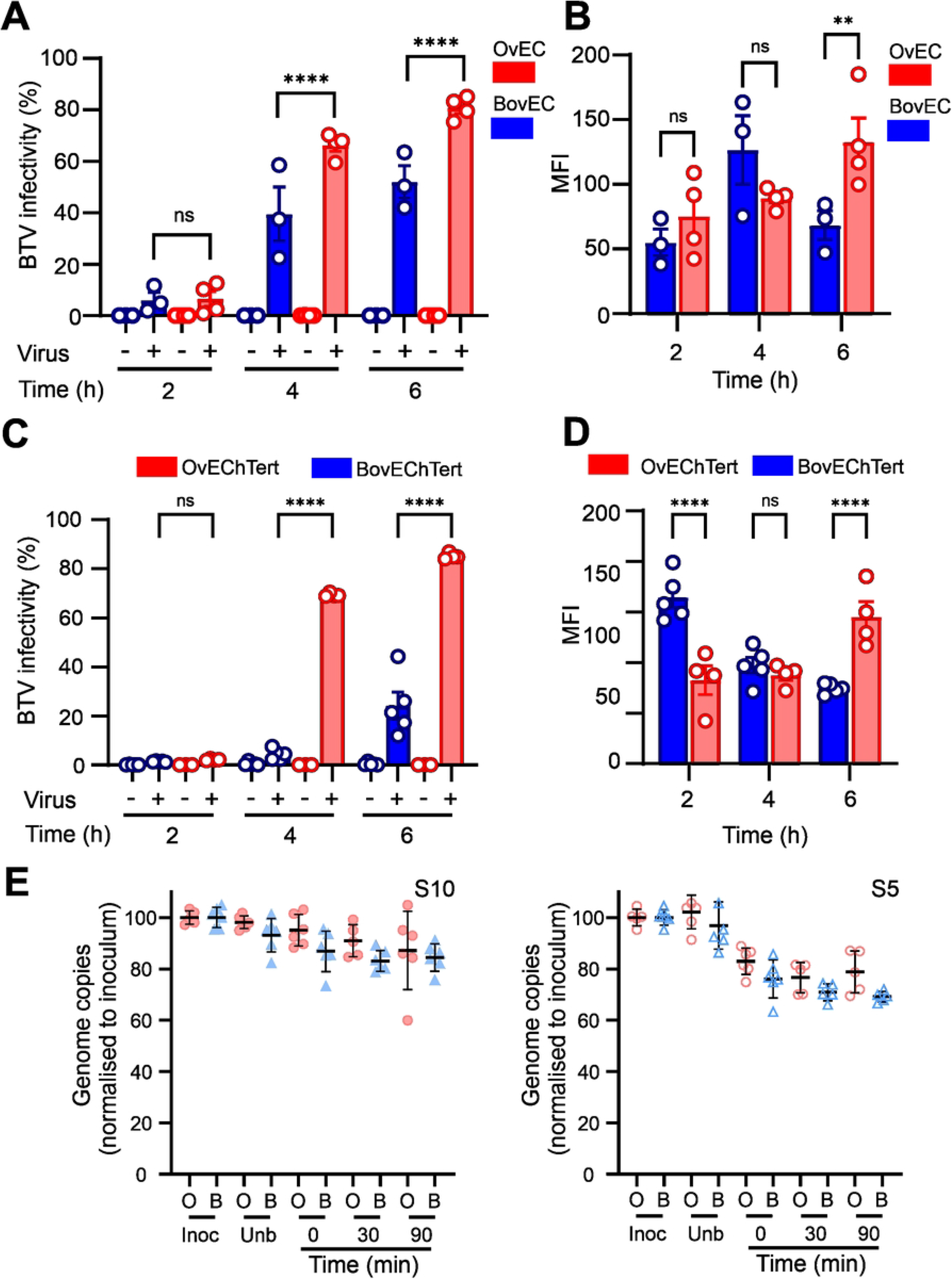
Relative restriction of BTV replication in bovine cells occurs within the first four hours post-infection. **(A).**Infectivity of BTV-8mGFP in primary OvEC and BovEC. Cells were infected with BTV8-mGFP at MOI∼1 and fixed at 2, 4 and 6 hpi. mGFP expression was determined by flow cytometry. Data was obtained from 4 independent experiments. **(B)**. Mean fluorescent intensity (MFI) of the infected primary OvEC and BovEC populations at different times post infection. Data from 4 independent experiments. **(C)** Infectivity of BTV-8mGFP in OvEChTert and BovEChTert cells. Experiments were carried out as in (A). **(D)** Mean fluorescent intensity (MFI) of the infected OvEChTert and BobEChTert. Data for (C) & (D) was obtained using the immortalised BovEC from 5 independent experiments using two different clones and data from immortalised OvEC is from 4 independent experiment using one clone (E). Binding and entry assays of BTV-8 into OvEChTert and BovEChTert cells assessed by qRT-PCR for either BTV segment 10 or 5 (S10; left panel and S5, right panel). Cells were synchronously infected BTV-8 at an MOI∼10 at 4°C, and samples taken from the inoculum, unbound virus, and wash. Cells harvested at 0-, 30- and 90-min post infection. Samples were normalised to the inoculum (set at 100%). Data is from 6 independent experiments.

We next tested whether BTV could enter OvEC more efficiently than BovEC. To this end, we infected immortalised OvEC and BovEC (MOI∼10) by spinoculation and harvest samples from the inoculum, the inoculum after spinoculation for 1h at 4°C (unbound fraction), as well as from cells at 0, 30 and 90 min post-infection. We then extracted and quantified viral RNA using qRT-PCR assays for either segment 5 or segment 10 and normalise values to the inoculum. We noticed a trend for more viral RNA to be bound and enter immortalised OvECs, although values were not significantly different (Fig 4E). Collectively, the data obtained so far suggest that the relative impairment of BTV replication in BovEC begins as early as 4 hpi.

### BTV-induced protein shutoff is more pronounced in ovine cells and downregulates ISG expression

Next, we investigated whether we could observe any species-specific differences in virus-induced host protein shutoff, a known mechanism used by BTV to modulate the cellular environment [63]. To this end, we infected OvECs and BovEC with BTV8 (using an MOI of 0.04 or 4), and then treated them with puromycin for 1h at 4, 16 or 24 hpi prior to harvest. Incorporation of puromycin into nascent peptides provides an indication of cellular protein synthesis [64]. As expected, BTV attenuated host protein synthesis both in OvEC and BovEC (Fig. 5A). However, when cells were infected with a high MOI, shutoff was essentially completed in OvECs by 16 hpi, while BovEC still supported nascent protein synthesis at 24 hpi (Fig 5A-B). We next examined BTV protein expression and NS1 was easily detectable in both OvEC and BovEC at 16 and 24 hpi (Fig 5A). We found a relatively higher expression of NS1 in OvEC as opposed to BovEC at low MOI (Fig 5C; 16 and 24 hpi p< 0.05 by 2-way ANOVA) while at high MOI the differences were not statistically significant (Fig 5C). In addition, we observed that both pSTAT1 and STAT1 were downregulated especially in OvEC (Fig 5A, D).

**Figure 5.**
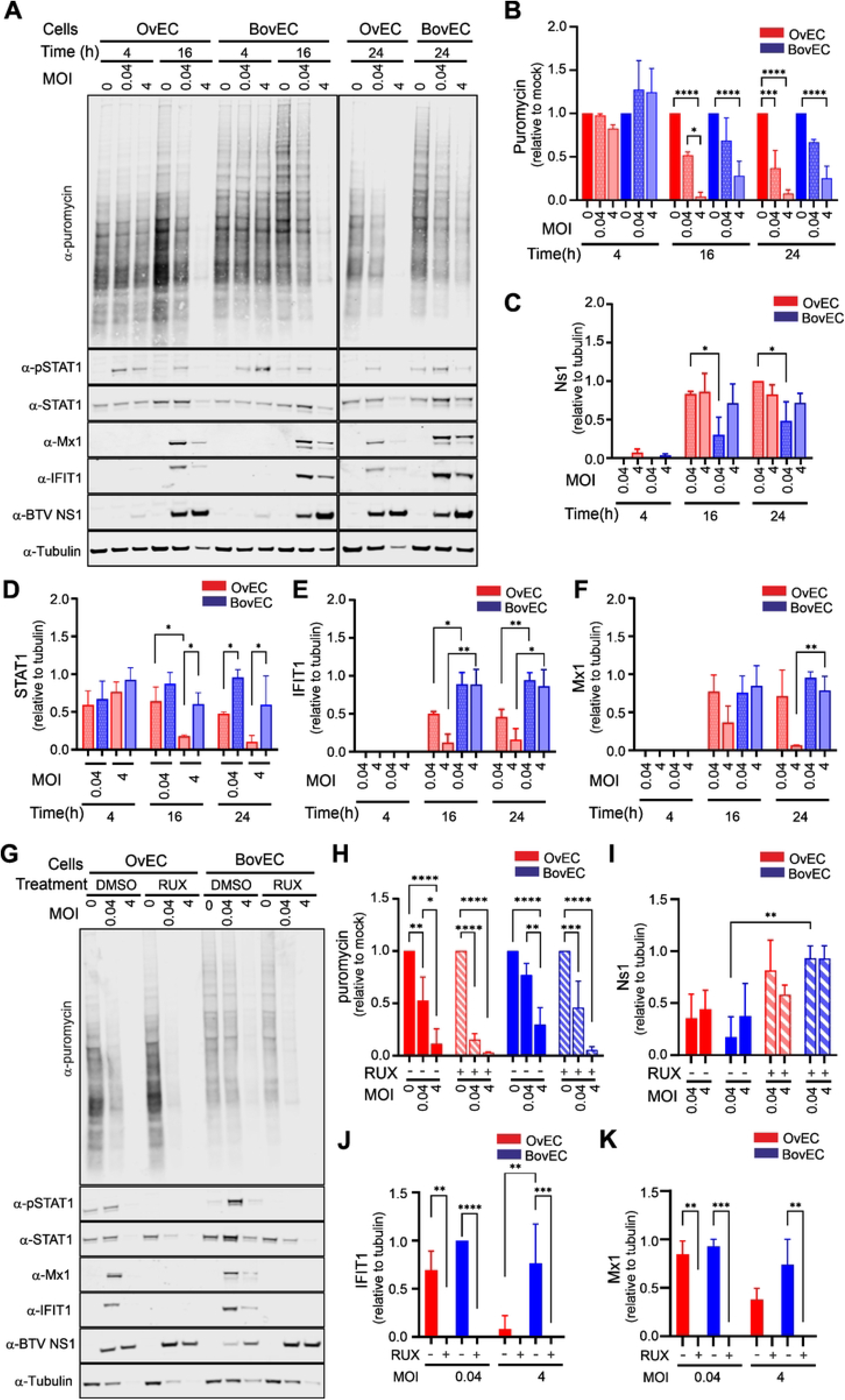
Virus induced global host cell protein shutoff by BTV is faster and more pronounced in ovine compared to bovine endothelial cells. **(A)**. Immunoblotting of phosphorylated STAT1, total STAT1, Mx1, IFIT1, BTV NS1 and protein translation rates (puromycin labelled treatment for 1 h) in BovEC and OvEC infected with BTV-8. Cells were infected at a 0.04 or 4 MOI and harvested at the timepoints indicated. (**B-F**) Relative quantification of immunoblots as in panel A obtained from 3 independent experiments. (B) Quantification of relative protein shutoff over the course of BTV infection as determine by puromycin incorporation. All values were normalised to the mock (set to 1) for each timepoint. (C). Relative expression of BTV NS1 over time normalised to tubulin in OvEC and BovEC infected with BTV-8. (D) Expression of STAT1 (relative to mock) over time normalised to tubulin in OvEC and BovEC infected with BTV-8. (E) Relative expression of IFIT1 over time normalised to tubulin in OvEC and BovEC. Cells were infected with BTV-8 using 2 different MOIs and harvested at 3 different timepoints. Data is from n=3 independent experiments. (F) Relative expression of Mx1 over time normalised to tubulin in OvEC and BovEC. Cells were infected with BTV-8 using 2 different MOIs and harvested at 3 different timepoints. Data is from n=3 independent experiments. **(G)**. Immunoblotting of phosphorylated STAT1, total STAT1, IFIT1, Mx1, BTV NS1 and protein translation rates (puromycin labelled treatment for 1 h) in OvEC and BovEC pre-treated with 4 μM RUX and infected with BTV-8. Cells were infected at a 0.04 or 4 MOI and harvested at 24 hpi. (**H-K**) Relative quantification of immunoblots as in panel G obtained from 3 independent experiments. (H) Relative protein shutoff in OvEC and BovEC infected with BTV as determine by puromycin incorporation. All values were normalised to the mock. (I) Relative expression of BTV NS1 (relative to tubulin) in OvEC and BovEC infected with BTV-8. (J) Relative expression of IFIT1 (normalised to tubulin) in OvEC and BovEC. (K) Relative expression of Mx1 in OvEC and BovEC. For panels H-K, cells were pre-treated with 4 μM Rux for 4 h prior to infection and was maintained in the media after infection. All data was from 3 independent experiments. Statistical significance was calculated using a Two-way ANOVA performed using Tukey’s multiple comparison test

During the IFN response, pSTAT1 is a key effector of the Jak/Stat signalling pathway leading to ISG upregulation. Both IFIT1 and Mx1 (as suitable ISG markers) were, as expected, upregulated in response to BTV infection (Fig 5A, E-F). However, when cells were infected at high MOI, downregulation of IFIT1 and Mx1 at late time-points was only apparent in ovine cells. Downregulation of ISGs and STAT1 were most likely a result of generalised host protein shutoff, as opposed to specific targeting by the virus, as the expression patterns displayed by IFIT1, Mx1 and STAT1 were very similar to those observed by the house-keeping gene α-tubulin (Fig. 5A). As mentioned above, α-tubulin was used as house-keeping gene control. Normally α-tubulin is used as gel loading control in each lane. However, in this series of experiments, because of virus-induced cellular shutoff, the intensity of the α-tubulin band was, as expected, significantly decreased in the samples where host protein synthesis was most downregulated. Hence, protein expression in each independent experiment was normalised to the average α-tubulin expression.

Similar experiments were also carried out in OvFib and BovFib (Figure S2). Virus-induced protein shutoff in fibroblasts was slower in both sheep and cattle fibroblasts compared to infected EC, with no statistical difference of this process in these cell lines. This is also in line with BovFib possessing a reduced ability to hamper BTV replication compared to BovEC.

We next assessed whether virus-induced protein shutoff is dependent or independent from the virus-induced IFN response. We repeated the experiments described above, focusing at 24 hpi, as differences between BTV infected sheep and cattle cells were most evident at this timepoint. We performed infections in the presence or Rux or DMSO (as a carrier control). Both Rux and DMSO were maintained in the media for the entire duration of the experiment. Host protein shutoff was more pronounced in cells treated with Rux, in both OvEC and BovEC, compared to those treated with DMSO (Fig 5G). Expression of BTV NS1 was, as expected increased, especially in BovEC, in cells treated with Rux compared to DMSO-treated controls (Fig 5H). Despite the Rux treatment, shutoff in BovEC was still not as pronounced as in OvEC, especially at low MOI (Fig 5H). As expected, the treatment with Rux prevented STAT1 phosphorylation and upregulation of ISGs (Fig 5G). As observed previously, in the presence of DMSO, the expression of Mx1 and IFIT1 was higher in BovEC infected with BTV8 (Fig 5J-K). These results indicate that the timing and extent of host protein shutoff is directly correlated to BTV replication levels. Blocking the IFN response by inhibiting the JAK/STAT pathway facilitates, virus-induced host shutoff. On the other hand, protein shutoff at later timepoints negatively affects STAT1 phosphorylation and therefore the IFN response.

### Higher magnitude of the IFN response to BTV infection in bovine cells

We next investigated whether we could identify in BTV-infected cells species-specific differences in the overall magnitude of the IFN response. We carried out transcriptomic analysis of endothelial cells infected at high MOI (∼10) with either BTV-2 or BTV-8 (using 5 biological replicates from different donors) and analysed them at 6 and 12 hpi.

To gain an overall understanding of the cell transcriptomic reaction to BTV infection, we first used the Reactome pathway enrichment analysis [65, 66]. At 6 hpi both in bovine and ovine cells infected with either BTV-2 or BTV-8, the top 5 most significantly enriched pathways were all related to cytokine and interferon signalling and the immune system (Fig. S3). At 12 hpi, other pathways unrelated to the immune response, such as metabolism of RNA, transcription and others were also significantly dysregulated in both bovine and ovine infected cells (Fig S3). Interestingly, when we carried out the same analysis using only those genes whose level of overexpression at 12 hpi was greater in infected bovine cells compared to sheep cells we obtained only four significant overlapping enriched pathways: (i) interferon signalling, (ii) cytokine signalling in immune system, (iii) interferon α/β signalling and (iv) immune system. Therefore, we next focused on analysing the transcriptional regulation of the type-I IFN system, (genes expressing the type-I IFN receptor, IFNAR1 and 2, and ISGs) in BTV infected cells.

In cultured cells, IFNβ is responsible for the first wave of the type-I IFN response as result of virus infection. Secreted IFNβ interacts with the type-I IFN receptor, which induces a signalling cascade resulting in the induction of ISGs and a second wave of both IFNβ and IFNα [67]. Type-I IFN genes in ruminants display numerous duplications and expansions, possessing at least 13 IFNα and 6 IFNβ paralogues [68, 69]. We concentrated only on those orthologues displaying non-zero expression during virus infection. Importantly, we observed no baseline differences in ISG expression, suggesting that baseline IFN expression, and downstream regulation is not different across species in the absence of infection in this model (Fig 6A). In both BTV-2 and BTV-8 infected animals, there was a noticeable upregulation of IFNβ genes in both sheep and cattle infected cells, at both 6 and 12 hpi (Fig 6B). At 6 hpi there seemed to be a trend for a higher upregulation of IFNβ genes in infected sheep cells but there were no differences at 12 hpi. For IFNα, levels of upregulation were minimal at 6 hpi while at 12 hpi we observed higher levels of upregulation in bovine infected cells (Fig 6C). We observed no major differences between sheep and cattle infected cells for the expression of type-I IFN receptors (Fig 6D). We next investigated whether we could identify any differences in the magnitude of ISG expression in infected sheep and cow cells. To model species-specific differences in ISG expression in cells infected with BTV-2 or BTV-8, we run four separate generalised linear models (GLMs) (using edgeR’s glmQLFit function) [70, 71] (2 for BTV2 and BTV8 at both 6 and 12 hpi). We focused our analysis on those ISGS that are present in both sheep and cattle as we previously published [60]. At 6 hpi, there were 801 and 473 ISGs that there were upregulated in BTV-2 and BTV-8 infected cells, respectively. Of these stimulated ISGs, only a relatively small number (∼3-4%) were more highly upregulated in cells from one species versus the other. At 12 hpi, approximately 900 ISGs were upregulated in both sheep and cow cells infected with either BTV-2 or BTV-8. However, in both cases there was a significant number of ISGs (26-40% of the total differentially expressed ISGs) that were more highly upregulated in infected cattle cells compared to sheep ones. Importantly, these differences are limited to ISGs as we noted no major differences when we carried out the same analyses with the whole transcriptome and exclude ISGs. Hence these data confirmed that at RNA level, BTV-infected cattle cells display a more robust IFN response compared to infected sheep cells.

**Figure 6.**
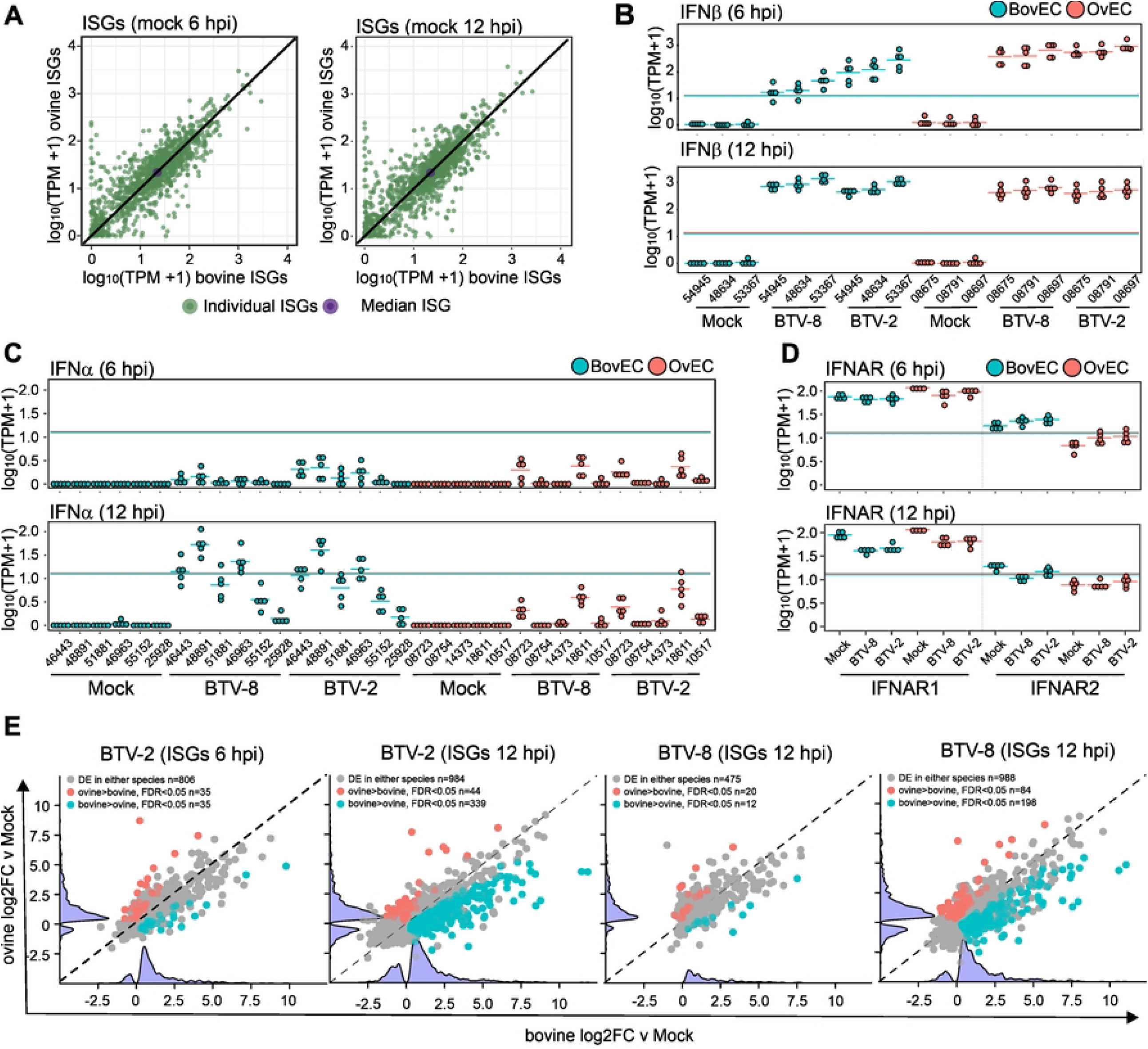
Transcriptomic response of BTV-infected OvEC and BovEC. (**A**) Scatter plot showing baseline regulation of ISGs in mock-infected OvEC and BovEC at 6 and 12hpi as identified by RNAseq. Values shown are transcripts per million (log10 (TPM+1)) and those identified in mock infected OvEC are plotted against the log10 (TPM+1) in mock infected BovEC. Individual ISGs are plotted green in and the median values of the ISGs plotted in purple. (**B**) Box-whisker plots of IFNβ transcripts in response to BTV-8 and BTV-2 infection in OvEC and BovEC at 6 & 12 hpi. (**C**) Box-whisker plots of IFNα transcripts in response to BTV-8 and BTV-2 infection in OvEC and BovEC at 6 & 12 hpi. (**D**) Box-whisker plots of IFNAR transcripts in response to BTV-8 & BTV-2 infection in OvEC and BovEC at 6 and 12 hpi. (**E**) Scatter plot showing differential expression (log2FC) of ISGs in OvEC and BovEC at 6 and 12 hpi in response to either BTV-2 or BTV-8 infection. Differentially expressed genes in both BovEC and OvEC are shown in grey, those only in OvEC in red and those in BovEC in blue. A violin plot of up- and down-regulated genes for each species is represented on each axis.

## DISCUSSION

Most animal reservoirs are tolerant to clinical disease, despite abundant virus replication that allows onward transmission to uninfected hosts [13]. However, the mechanisms underpinning disease tolerance in reservoir species have not been investigated as widely as those occurring during disease pathogenesis in susceptible hosts. Understanding the complex virus-host interactions governing disease tolerance can allow us to interpret the critical balance between protective and damaging host responses [3]. The host innate immune responses, and in particular the type-I interferon (IFN) as one of its key effectors, plays a critical role in this balance as it stimulates the expression of antiviral proteins, leading to host resistance, but at the same time can also exacerbate tissue lesions and disease severity through the release of pro-inflammatory factors [60, 72].

In this study, we used BTV, a virus of ruminants, as a model system to investigate the complex virus-host interactions in susceptible and tolerant species. BTV infects a wide range of ruminants resulting in drastically different clinical manifestations: whilst sheep in general develop clinical signs and can succumb to the disease, cattle remain mostly asymptomatic despite showing long lasting viremia and being a reservoir of infection [25, 26, 38]. We showed that the BTV replication kinetics are different whether infections occur in cells derived from sheep or cattle. We used both primary skin fibroblasts and endothelial cells in our study, the latter being the key target of BTV replication and pathogenesis *in vivo* [37, 39]. The phenotype observed was similar in both cell types, although more pronounced in ECs. We observed some variation between cell types and virus serotypes, but we accounted for the inherent variation of different primary cell preparations and the diverse genetic background, by using distinct batches of cells from multiple donors, and repeating the experiments independently several times. In addition, we used EC within 3 passages from isolation, with no freeze-thaw cycles in between.

We determined that as early as 4 hpi, the replication kinetics of BTV were relatively hampered in cattle cells, compared to sheep cells. Hence, the very early events of BTV replication, are more efficient in sheep cells. Using two distinct qRT-PCR assays to quantify viral RNA collected at 0, 30 and 90 min post-infection, we observed a trend showing higher levels of BTV RNA in infected sheep cells compared to cattle cells, although these differences did not reach statistical significance.

Most importantly, we established that the differences in BTV replication kinetics in cattle and sheep cells were largely abrogated by inhibiting the Jak-Stat pathway, whose activation leads to the upregulation of hundreds of interferon stimulated genes (ISGs) [60]. In addition, pre-treatment with IFN had a relatively minor effect on the replication kinetics of BTV in sheep cells while it substantially decreased viral replication in cattle cells.

Hence, BTV shows a delayed replication kinetics in cattle cells that is largely abrogated by the inhibition of the IFN response. We identified in this study several ISGs with anti-BTV properties: IFIT1, RSAD2, OAS2, UBA7 and HAS3. For both IFIT-1 and RSAD-2, we were able to confirm their anti-BTV activity both in overexpression and gene knock-out experiments. IFIT1 and RSAD2 have been shown before to possess antiviral activity for a wide variety of viruses. IFIT-1 acts as a repressor of mRNA translation by interacting and interfering with the translation initiation factor eIF3 and acting on transcripts with the 2’O-methylation of the 5’ RNA cap structure [73]. RSAD2 (encoding viperin) catalyses the conversion of cytidine triphosphate (CTP) to 3’-deoxy-3’,4’-didehydro-CTP (ddhCTP), causing premature termination of RNA synthesis by the polymerase of various RNA viruses [74]. However, we found no differences in the antiviral properties of either the IFIT-1 and RSAD-2 orthologues, or of any of the other anti-BTV ISGs identified in our study. These data therefore suggest that the effectors of both the sheep and cattle IFN response are equally equipped to hamper the replication kinetics of BTV.

To overcome the innate immune response, many viruses, including BTV, employ host protein shutoff as a mechanism to suppress the translation of antiviral proteins [75-79]. Indeed, we showed that protein shutoff was more pronounced and occurred at earlier time points after infection of sheep compared to cattle cells. As expected, host protein shutoff included pStat1 (a phosphorylated protein essential for ISG expression) and antiviral ISGs such as MX1 and IFIT1, whose expression was relatively lower in sheep cells compared to cattle cells. Critically, host protein shutoff was not dependent on the IFN response as it occurred also in the presence of an inhibitor of the JAK/STAT signalling. On the contrary, host shutoff was more pronounced in cells treated with Ruxolitinib, providing further evidence that this process is directly linked to the levels of BTV proteins expression. Indeed, BTV-induced protein shutoff and modulation of the type-I IFN response has been shown to be linked to expression of viral proteins NS1, NS3 and NS4 [78-81].

Using RNAseq in BovEC and OvECs infected with a high MOI, we found no major differences in the induction of IFN-β or the type-I IFN receptor (IFNAR1/2). Indeed, ISG upregulation as a result of BTV infection was relatively similar in sheep and cattle cells at early time point after infection. However, at later time points (12 hpi), a higher number of ISGs were upregulated in cattle compared to sheep cells. We also observed higher levels of upregulation of IFNα at 12 hpi.

Hence, our data collectively show that relatively small differences in viral replication determine a more efficient modulation of the type-I IFN response in cells of tolerant hosts, as opposed to susceptible ones. The head start of BTV replication in sheep cells results in a more abundant expression of viral proteins at earlier timepoints after infection, resulting in a faster virus-induced host protein synthesis shutoff compared to infected cattle cells. BTV modulates the cell IFN response more effectively in sheep cells, as a direct result of (i) virus-induced inhibition of translation of antiviral proteins and likely (ii) of any of the cellular proteins involved in the signalling pathways leading to ISG upregulation. In addition, earlier expression of BTV proteins in infected sheep cells could also lead to an earlier modulation of the IFN response as a direct effect of the expression of IFN antagonists such as NS3 and NS4 [79-82]. Conversely, in BTV-infected cattle cells, a relatively lower expression of viral proteins at earlier timepoints allows the IFN response to be activated at earlier timepoints and slow down viral replication. Importantly, the data obtained *in vitro* are compatible with previously published in vivo observations, showing a delayed onset of viremia in experimentally infected cattle compared to sheep [42, 97-99].

Our study suggests that the relative tipping point between tolerant and susceptible hosts during viral infection can be likely determined by small differences in the modulation of the IFN response. The innate immune response, and in particular the type-I IFN response, are described as the first line of antiviral responses available to the host. However, in reservoir species, the balance between virus and innate immunity has been finely tuned through evolution to favour both host and virus survival through evolution. Counterintuitively, we propose a model where in reservoir species the timing and extent of the type-I IFN response ultimately facilitates virus transmission from infected to uninfected hosts.

Overall, it will be extremely important to determine the mechanisms underpinning disease tolerance not only to understand important lessons in virus emergence and pathogenesis but also to provide new intellectual frameworks for the development of host-directed antiviral therapeutic strategies [6].

## MATERIALS AND METHODS

### Cells

BSR and CPT-Tert cells were maintained as previously described [80, 83]. *Ex vivo* samples from healthy cattle and sheep (ears and aortas) were harvested from abattoirs and used to isolate primary fibroblasts and endothelial cells following the previously published protocols [60, 84]. All isolated cells were tested for pestiviruses and mycoplasma prior to use, and passages of the primary cells were kept to a minimum. Primary cells used in this study are referred from now on as OvEC (ovine endothelial cells), OvFib (ovine fibroblasts), BovEC (bovine endothelial cells) and BovFib (bovine fibroblasts). We took care to use at multiple donors to prepare primary cells.

Bovine and ovine primary cells were also immortalised by transducing cells with a retrovirus expressing the catalytic subunit of human telomerase (hTERT), as previously described [85]. Briefly, cells were immortalised using the catalytic subunit of hTERT delivered by transduction using a lentivirus vector (pLV-hTERT-IRES-Hygro; Addgene). Two days post-transduction, cells were selected with 200μg/ml of hygromycin for 7 days and maintained as a bulk population for fibroblasts. Immortalised cell lines generated in this study are referred to a BovEChTert, OvEChTert, BovFibTt and OvFibT. Morphology, growth rate as well as IFN response of immortalised fibroblasts and endothelial cells were monitored over time (up to passage 15) and were similar to relative bovine and ovine primary cells. As previously described, this system has been demonstrated to extend the life span of the cells without compromising interferon signalling [85, 86]. Single cell clones of endothelial cells were selected, amplified, and screened for interferon competency.

In some experiments, as indicated below, cells were treated with the following drugs. Ruxolitinib (Rux; APEXBio) a potent inhibitor of the Janus kinase (JAK)-signal transducer and activator of transcription (STAT) pathway; universal interferon (uIFN; PBL InterferonSource); puromycin dihydrochloride (Sigma Aldrich).

### Viruses

Two strains of BTV were used in this study, BTV-8 and BTV-2 as previously described [32, 80]. A BTV-8mGFP virus expressing a monomeric GFP in the C-terminus of the NS1 open reading frame was generated by reverse genetics using established methods [80, 87, 88]. Upon rescue, BTV-8mGFP was plaque purified for the generation of working stocks. BTV-8mGFP maintained GFP expression for at least two passages, and aliquots of passage 2 were stored at −70°C and were used to generate the working stock used in all the experiments.

The replication kinetics of BTV were determined in ovine and bovine primary cells. Cell monolayers in a 12-well plate were infected at a MOI of 0.01 in different conditions. To inhibit the JAK/STAT pathway, cells were pre-treated for 4 h with 4 μM Rux or mock-treated with the equivalent volume of DMSO diluent. Cells were then maintained in media supplemented with Rux for the duration of the experiments. To stimulate an antiviral state cell were pre-treated with 500-1000 IU/mL of uIFN for 24 h prior to infection and uIFN was supplemented into the media for the duration of the experiment.

At different timepoints post-infection, 100 μL of supernatant was collected from each well and replaced with 100 μL of the appropriate cell media. Supernatants were stored at 4°C until they were processed for titration. Virus titres were determined by end-point dilution analysis on BSR cells and expressed as log_10_ TCID_50_/mL [89, 90]. Each virus growth curve experiment was performed at least 3 times independently and each experiment was carried out in triplicate. At least two independently generated virus stocks were used for each virus.

### Visualisation of cytopathic assay in primary cells

Primary ovine and bovine endothelial cells were infected in 12 well-plates as described previously. Infected cells were incubated at 37°C and fixed in 4% (w/v) formaldehyde in PBS (FS) at 24, 48, 72 and 96 hpi. Monolayer were stained with Coomassie Blue staining solution (0.1%[w/v] Coomassie Brilliant Blue R-250; 45%[v/v] methanol; 10%[v/v] glacial acetic acid) and imaged with a photo scanner (Epson Expression 1680 Pro).

### Bovine interferon stimulated genes (ISG) library

A bovine arrayed ISG library was synthesised (Genewiz) in the pSCRPSY lentiviral vector allowing the co-expression of the gene of interest and a fluorescent protein (TagRFP) as previously described for human and macaque ISG library [56, 61]). In total, 289 unique bovine genes and 25 additional cDNA sequences accounting for splicing variants were included in the library (Supplementary Table 1). They correspond to the repertoire of the most positively up-regulated genes identified in bovine primary cells following IFN treatment [60] in addition to the mammalian “core ISGs” evolutionarily conserved across mammalian and chicken genomes. Lentivirus vectors expressing the ISG library were generated in 96-well plate format as described previously [56], with one ISG per well, by co-transfecting 3.5×10^4^ adherent 293T cells with 125 ng of the ISG construct in the SCRPY vector and plasmids expressing HIV gag-pol (NLGP) and the vesicular stomatitis virus glycoprotein (VSV-G) in a ratio of 25:5:1, respectively. Media was changed 24h post-transfection and supernatants were subsequently harvested at 48, 72, and 96 h, and stored at −80°C. Independent stocks of lentiviruses were generated for validation assays.

### Bovine ISG screens

293T suspension cells were transduced via spinoculation for 1h at 500 x *g* at 4°C in the presence of 8 μg/mL polybrene. The percentage of RFP-positive cells is indicative of the transduction efficiency and was used as an indicator of ISG expression levels. Forty-eight hours post transduction, the cell populations were split and infected with 50 μL of BTV8-mGFP virus for a period of 16 h or 32 h. The dose of BTV-8mGFP had previously been optimised to ensure between 10 and 50% of cells in each well were infected. For the late screen, a final concentration of 0.625 μg/mL of puromycin was supplemented in the media, enabling virus replication to be limited to the transduced cells. At 16 and 32 hpi, cells were washed, trypsinised and fixed in 4% (w/v) formaldehyde in PBS. Cells were then analysed by flow cytometry, using the Guava Easycyte instrument (Millipore) equipped with a 488 nm and a 532 nm laser. For each well, single live cell population was gated and levels of GFP and RFP were measured. ISG with transduction levels (measured by RFP expression) below 10% are excluded from downstream analyses. Data was then analysed using the FlowJo (Treestar) software.

### Western blotting

Cells were lysed as described previously [91]. Samples were subjected to SDS-PAGE and immunoblotting was performed using the following antibodies: puromycin (Millipore, MABE343), pSTAT1 (Cell Signaling, 9167S), STAT1 (Santa Cruz, sc-592), IFIT1 (Origene, TA500948), BTV NS1 [80], RSAD2 (Proteintech, 11833-1-AP), α-tubulin (Proteintech, 66031-1-Ig) and GAPDH (Cell Signaling, 2118S). Mx1 antibody was kindly provided by Dr. Georg Kochs from University Medical Centre, Freiburg Germany. The secondary antibodies used for immunoblotting were anti-rabbit IgG (H+L) (DyLight™ 800 4X PEG Conjugate) (Cell Signaling, 5151S) and anti-mouse IgG (H+L) (DyLight™ 680 Conjugate) (Cell Signalling, 5470S). Proteins were visualized using the Odyssey CLx imager and quantification was performed using the Image Studio Lite software (LI-COR Biosciences).

### CRISPR/Cas9 knock-out cells

Target-specific single guide RNAs (sgRNAs) for bovine IFIT1 and RSAD2 (*IFIT1* forward, 5’ caccgAATCGTTGTCTATCGCCTGG 3’ and reverse 5’ aaacCCAGGCGATAGACAACGATTc 3’; *RSAD2* forward 5’ caccgAGTGGTAATTGACGCTGGTG 3’ and reverse 5’ aaacCACCAGCGTCAATTACCACTc 3’) were designed using CHOPCHOP web tool (https://chopchop.cbu.uib.no/) and cloned into pLentiCRISPR v2 vector (Addgene) as described [92, 93]. Lentivirus were made as described above and used to transduce immortalised bovine cells (BovEChTert and BovFibhTert). Cells were selected in the presence of 2.5 mg/ml puromycin, and single cell population expanded. To ensure that resulting cells maintained their responsiveness to interferon and the gene of interest had been knocked out, cells were treated with 500-1000 U uIFN for 24h at 37°C and processed for immunoblotting for relevant proteins. For each ISG, two clones with the greatest reduction in specific protein expression following uIFN treatment were defined as knockout (KO) bovine fibroblasts and used for further experimentation.

### BTV-mGFP infectivity in primary bovine and ovine endothelial cells

BovEC and OvEC from aortas collected from three different animals were seeded in 96-well plates. Two-fold serial dilutions of BTV8-mGFP were used to spinoculate cells at 500 x *g* for 1 h at 4°C. The inoculum was then removed, cells were washed with serum free medium once before adding warm endothelial cell growth medium. Infected cells were then incubated at 37°C for 6 h and then fixed in 4% (w/v) FS. The proportion of GFP-expressing cells was then determined by flow cytometry (Guava® easyCyte, Luminex) by acquiring at least 10,000 events. In addition, OvECs, BovECs, and immortalised OvEC-hTert and BovECs-hTert were infected with BTV-mGFP (MOI of either 0.3 or 1) and fixed at either 2, 4 or 6 hpi. Percentage of GFP-expressing cells, in addition to the mean fluorescence intensity (MFI) was obtained by flow cytometry.

### Entry assays

OvEC-hTert and BovECs-hTert cells were seeded in a 12 well-plate and at confluency were infected with an MOI of 1 synchronously at 4°C by spinoculation as described above. The inoculum was aspirated, and cells washed in ice with cold media twice to remove unbound virus. Aliquots of the wash were kept for RNA extraction. Media (pre-warmed to 37°C) was added directly to the cells which were further incubated at 37°C. At the specified time post infection media was collected and Trizol® (Invitrogen) was directly added to the monolayer. RNA was extracted from the samples as described before [60] and the number of BTV viral genome equivalent determined by quantitative RT-PCR based on number of copies of BTV segment 5 and segment 10 as described before [94]. All reactions were performed on ABI 7500 Fast thermal cycler.

### Protein shutoff assays

To examine protein shutoff nascent proteins were labelled with puromycin as previously described [84]. Primary ovine and bovine cells were infected with BTV-8 at either MOI of 0.04 or 4 by spinoculation as described above. After 4, 16 and 24 hpi, puromycin was added to the cells at a final concentration of 3.3 ng/ml for 1 h. Following cell lysis, samples were subjected to SDS-PAGE and immunoblotting was performed for puromycin and various other proteins as indicated. For experiments in the presence of Rux, cells were pre-treated with 4 μM Rux for 4 h prior to spinoculation at 500 x *g* and Rux-supplemented media (or the DMSO control) was added back onto the cells for the duration of the experiment. At 24 hpi, 3.3 ng/ml puromycin was added to the cells for 1 h. Cells were then lysed and processed for immunoblotting as mentioned above.

Relative quantification of puromycin, Mx1, IFIT1, STAT1 and NS1 was performed by normalizing against average α-tubulin for each condition to compensate for the reduction in expression of the housekeeping genes due to protein shutoff. For the quantification of the puromycin labelling in infected samples, the value of each sample was normalized against mock infected cells (set to 1). Relative quantification of other viral and cellular proteins mentioned above was instead performed using as unit the highest value (set to 1) for each MOI in each experiment.

### RNAseq

Primary ECs were infected in 12-well plates at MOI∼5 with either BTV-8 or BTV-2 by spinoculation at 500 x *g* at 4°C for 1 h. Cells were then further incubated 1 h at 4°C. Media was then removed and cells were washed with warm media before incubation at 37°C for either 6 or 12 hpi. Monolayers were then washed with warm PBS to remove any cellular debris and immediately lysed in 750 μL of Trizol. Total RNA was extracted from each sample as previously published [60]. Sample RNA concentration was measured with a Qubit Fluorimeter (Life Technologies) and the RNA integrity was determined using an Agilent 4200 TapeStation. Samples had an average RNA Integrity Number of ∼9.8 and 500 ng of total RNA from each sample was taken for library preparation. Libraries for sequencing were generated using an Illumina TruSeq Stranded mRNA HT kit, according to the manufacturer’s instructions. Briefly, polyadenylated RNA molecules were captured, followed by fragmentation. RNA fragments were reverse transcribed and converted to dsDNA, end repaired, A-tailed, ligated to indexed adaptors and PCR amplified. Libraries were pooled in equimolar concentrations and sequenced in an Illumina NextSeq 500 and 550 sequencers using a high output cartridge, generating single reads with a length of 75 bp. At least 94.77% of the reads generated presented a quality score of 30 or above.

RNAseq read quality was assessed using the FastQC software, and sequence adaptors were removed using TrimGalore. The reference Ovis aries genome (Oar_v3.1) and the Bos taurus genome (ARS-UCD1.2) were downloaded from the Ensembl genome browser and RNAseq reads were then aligned to their respective host genomes. Following the alignment, FeatureCount [95] was used to count reads mapping to genes in annotation files. The EdgeR package was then used to calculate the gene expression levels and to analyse differentially expressed genes [71]. The raw FASTQ files generated during this project have been submitted to the European Nucleotide Archive (ENA) under project accession number PRJEB55791.

To model the species-specific responses to infection with two different BTV strains, four separate GLMs were run using edgeR’s glmQLFit function: 2 for BTV2 and BTV8 at both 6 and 12 h post infection. For each GLM, the five replicate read counts from each of the four conditions (bovine, ovine, infected, and uninfected) were used, giving 5×4 sets of counts input for each GLM. These counts were used to generate four effect terms. Each effect term describes the log2 fold change (log2FC) between the conditions, with an associated FDR value for each term to test for statistical significance of the effect: (i) An uninfected sheep cell term, describing baseline expression in uninfected cells, a term for cow expression relative to sheep; (ii) a cow term providing the cow-specific baseline expression in the cell lines; (iii) an infection term describing the logFC between uninfected and infected sheep; and (iv) a cow-by-infection interaction term describing the difference in the way the two species’ cell lines respond to infection. Partitioning these effects allowed the species-specific response to infection to be determined thus avoiding the confounding effects of baseline expression that would be involved in a simple pairwise comparison across species. The fact each effect is associated with an FDR value contextualises the effect size.

A model matrix for each GLM was also calculated. An intercept was calculated to provide the baseline value for bulk uninfected sheep cell sequencing. It was arbitrary if ovine or bovine data provided the baseline intercept, it just inverted the second and fourth species-specific effect terms. Transcripts per million are calculated using the following equation: 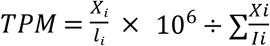. To gain an overall understanding of the cell transcriptomic reaction to BTV infection, we used the Reactome pathway enrichment analysis suite on each species’ response to infection [65, 66, 96]. We mapped ovine Ensembl IDs to human entrez IDs for this package using the biomaRt package in R[97].

**Statistical analyses** were performed using the GraphPad Prism software. Quantitative data were presented as arithmetic mean ± SEM. Statistical significance of differences in puromycin labelling between mock and MOI 0.04/MOI 4 at different time points was assessed using unpaired t-tests with Bonferroni post-test to compare each MOI with the mock infected sample. The same method was used to assess differences in NS1, Mx1, IFIT1 and RSAD2 expression between ovine and bovine samples at different time points. Differences in means were considered significant if P < 0.05; *P < 0.05, **P < 0.01, ***P < 0.001.

## Acknowledgements

This study was funded by an Investigator Award from the Wellcome Trust (206369/Z/17/Z). Additional funding was provided by the MRC (MC_UU_12014/10; MC_UU_12014/12). Authors contribution is the following. AH: conceptualisation, methodology, investigation; writing - review and editing; SB: methodology, investigation, formal analysis, writing - review and editing; WF: methodology, investigation, formal analysis, writing - review and editing; OM: methodology, investigation, software, formal analysis, data curation, visualisation, writing - review and editing; QG: methodology, investigation, software, formal analysis, data curation; MVarj: methodology, investigation, writing - review and editing; MV: methodology, investigation; MAA: methodology, investigation; AS; methodology, writing - review and editing; RMP: methodology; NCR: methodology; investigation; CM: methodology, resources; AT: methodology, AF: methodology, investigation; RER: supervision; SW: conceptualisation, methodology, supervision, resources, writing – review and editing; MES: methodology, investigation, formal analysis, visualisation, supervision, writing – original draft; MP: conceptualisation, supervision, project administration, funding acquisition, writing – original draft.

## Supporting Information

**S1 Fig. Cytopathic effect induced by BTV-8 in ovine and bovine primary aorta endothelial cells**. Cells were infected with BTV-8 at an MOI∼0.01 and monolayer fixed at different times points post infection. Monolayer integrity was assessed by staining cells with Coomassie brilliant blue.

**S2 Fig. Virus induced global host cell protein shutoff by BTV in ovine and bovine primary fibroblasts. (A)**. Immunoblotting of phosphorylated STAT1, total STAT1, Mx1, IFIT1, BTV NS1 and protein translation rates (puromycin labelled treatment for 1 h) in OvFib and BovFib with BTV-8. Cells were infected at a 0.04 or 4 MOI and harvested at the timepoints indicated. (**B-F**) Relative quantification of immunoblots as in panel A obtained from 3 independent experiments. **(B)** Quantification of relative protein shutoff over the course of BTV infection as determine by puromycin incorporation. All values were normalised to the mock (set to 1) for each timepoint. (C). Relative expression of BTV NS1 over time normalised to tubulin in OvFib and BovFib infected with BTV-8. (D) Expression of STAT1 (relative to mock) over time normalised to tubulin in OvFib and BovFib infected with BTV-8. (E) Relative expression of IFIT1 over time normalised to tubulin in OvFib and BovFib. Cells were infected with BTV-8 using 2 different MOIs and harvested at 3 different timepoints. Data is from n=3 independent experiments. (F) Relative expression of Mx1 over time normalised to tubulin in OvFib and BovFib. Cells were infected with BTV-8 using 2 different MOIs and harvested at 3 different timepoints. Data is from n=3 independent experiments.

**S3 Fig. Reactome pathway analysis of mapped differentially expressed genes in BTV infected bovine and ovine endothelial cells**. Pathways are indicated at both 6 and 12 hours post infection for cells infected with either BTV-2 or BTV-8. The −log10(Q value) for the significant pathways identified are plotted.

**S1 Data. Data of the bovine ISG over-expression screens**. Data includes gene names (including ENSEMBL ID), percentage of cells successfully transduced by each gene and the value of BTV-8 infectivity normalised to the mean value obtained across the entire library.

## REFERENCES

1. Haydon DT, Cleaveland S, Taylor LH, Laurenson MK. Identifying reservoirs of infection: a conceptual and practical challenge. Emerg Infect Dis. 2002;8(12):1468–73. Epub 2002/12/25. doi: 10.3201/eid0812.010317. PubMed PMID: 12498665; PubMed Central PMCID: PMCPMC2738515.

2. Casadevall A, Pirofski LA. Host-pathogen interactions: basic concepts of microbial commensalism, colonization, infection, and disease. Infect Immun. 2000;68(12):6511–8. Epub 2000/11/18. doi: 10.1128/IAI.68.12.6511-6518.2000. PubMed PMID: 11083759; PubMed Central PMCID: PMCPMC97744.

3. Rouse BT, Sehrawat S. Immunity and immunopathology to viruses: what decides the outcome? Nat Rev Immunol. 2010;10(7):514–26. Epub 2010/06/26. doi: 10.1038/nri2802. PubMed PMID: 20577268; PubMed Central PMCID: PMCPMC3899649.

4. McCarville JL, Ayres JS. Disease tolerance: concept and mechanisms. Curr Opin Immunol. 2018;50:88–93. Epub 2017/12/19. doi: 10.1016/j.coi.2017.12.003. PubMed PMID: 29253642; PubMed Central PMCID: PMCPMC5884632.

5. Medzhitov R, Schneider DS, Soares MP. Disease tolerance as a defense strategy. Science. 2012;335(6071):936–41. Epub 2012/03/01. doi: 10.1126/science.1214935. PubMed PMID: 22363001; PubMed Central PMCID: PMCPMC3564547.

6. Soares MP, Teixeira L, Moita LF. Disease tolerance and immunity in host protection against infection. Nature reviews Immunology. 2017;17(2):83–96. Epub 2017/01/04. doi: 10.1038/nri.2016.136. PubMed PMID: 28044057.

7. Schneider DS, Ayres JS. Two ways to survive infection: what resistance and tolerance can teach us about treating infectious diseases. Nature reviews Immunology. 2008;8(11):889–95. Epub 2008/10/18. doi: 10.1038/nri2432. PubMed PMID: 18927577; PubMed Central PMCID: PMCPMC4368196.

8. Raberg L, Graham AL, Read AF. Decomposing health: tolerance and resistance to parasites in animals. Philos Trans R Soc Lond B Biol Sci. 2009;364(1513):37–49. Epub 2008/10/18. doi: 10.1098/rstb.2008.0184. PubMed PMID: 18926971; PubMed Central PMCID: PMCPMC2666700.

9. Roy BA, Kirchner JW. Evolutionary dynamics of pathogen resistance and tolerance. Evolution. 2000;54(1):51–63. Epub 2000/08/11. doi: 10.1111/j.0014-3820.2000.tb00007.x. PubMed PMID: 10937183.

10. Raberg L, Sim D, Read AF. Disentangling genetic variation for resistance and tolerance to infectious diseases in animals. Science. 2007;318(5851):812–4. Epub 2007/11/03. doi: 10.1126/science.1148526. PubMed PMID: 17975068.

11. Read AF, Graham AL, Raberg L. Animal defenses against infectious agents: is damage control more important than pathogen control. PLoS Biol. 2008;6(12):e4. Epub 2009/02/19. doi: 10.1371/journal.pbio.1000004. PubMed PMID: 19222305; PubMed Central PMCID: PMCPMC2605932.

12. Regoes RR, McLaren PJ, Battegay M, Bernasconi E, Calmy A, Günthard HF, et al. Disentangling human tolerance and resistance against HIV. PLoS Biol. 2014;12(9):e1001951. Epub 2014/09/17. doi: 10.1371/journal.pbio.1001951. PubMed PMID: 25226169; PubMed Central PMCID: PMCPMC4165755.

13. Mandl JN, Ahmed R, Barreiro LB, Daszak P, Epstein JH, Virgin HW, et al. Reservoir host immune responses to emerging zoonotic viruses. Cell. 2015;160(1-2):20–35. Epub 2014/12/24. doi: 10.1016/j.cell.2014.12.003. PubMed PMID: 25533784; PubMed Central PMCID: PMCPMC4390999.

14. Barman TK, Metzger DW. Disease Tolerance during Viral-Bacterial Co-Infections. Viruses. 2021;13(12). Epub 2021/12/29. doi: 10.3390/v13122362. PubMed PMID: 34960631; PubMed Central PMCID: PMCPMC8706933.

15. Irving AT, Ahn M, Goh G, Anderson DE, Wang LF. Lessons from the host defences of bats, a unique viral reservoir. Nature. 2021;589(7842):363–70. Epub 2021/01/22. doi: 10.1038/s41586-020-03128-0. PubMed PMID: 33473223.

16. Oura CA, Powell PP, Anderson E, Parkhouse RM. The pathogenesis of African swine fever in the resistant bushpig. J Gen Virol. 1998;79 (Pt 6):1439–43. Epub 1998/06/20. doi: 10.1099/0022-1317-79-6-1439. PubMed PMID: 9634086.

17. Sanchez-Cordon PJ, Nunez A, Neimanis A, Wikstrom-Lassa E, Montoya M, Crooke H, et al. African Swine Fever: Disease Dynamics in Wild Boar Experimentally Infected with ASFV Isolates Belonging to Genotype I and II. Viruses. 2019;11(9). Epub 2019/09/22. doi: 10.3390/v11090852. PubMed PMID: 31540341; PubMed Central PMCID: PMCPMC6783972.

18. Gethmann J, Probst C, Conraths FJ. Economic Impact of a Bluetongue Serotype 8 Epidemic in Germany. Front Vet Sci. 2020;7:65. Epub 2020/03/03. doi: 10.3389/fvets.2020.00065. PubMed PMID: 32118078; PubMed Central PMCID: PMCPMC7034324.

19. Hasler B, Howe KS, Di Labio E, Schwermer H, Stark KD. Economic evaluation of the surveillance and intervention programme for bluetongue virus serotype 8 in Switzerland. Prev Vet Med. 2012;103(2-3):93–111. Epub 2011/10/25. doi: 10.1016/j.prevetmed.2011.09.013. PubMed PMID: 22018548.

20. Rushton J, Lyons N. Economic impact of Bluetongue: a review of the effects on production. Vet Ital. 2015;51(4):401–6. Epub 2016/01/08. doi: 10.12834/VetIt.646.3183.1. PubMed PMID: 26741252.

21. Tabachnick WJ. Culicoides variipennis and bluetongue-virus epidemiology in the United States. Annual review of entomology. 1996;41:23–43. Epub 1996/01/01. doi: 10.1146/annurev.en.41.010196.000323. PubMed PMID: 8546447.

22. Belbis G, Zientara S, Breard E, Sailleau C, Caignard G, Vitour D, et al. Bluetongue Virus: From BTV-1 to BTV-27. Adv Virus Res. 2017;99:161–97. Epub 2017/10/17. doi: 10.1016/bs.aivir.2017.08.003. PubMed PMID: 29029725.

23. Huismans H, Erasmus BJ. Identification of the serotype-specific and group-specific antigens of bluetongue virus. Onderstepoort J Vet Res. 1981;48(2):51–8. Epub 1981/06/01. PubMed PMID: 6273773.

24. Ries C, Sharav T, Tseren-Ochir EO, Beer M, Hoffmann B. Putative Novel Serotypes ‘33’ and ‘35’ in Clinically Healthy Small Ruminants in Mongolia Expand the Group of Atypical BTV. Viruses. 2020;13(1). Epub 2021/01/02. doi: 10.3390/v13010042. PubMed PMID: 33383902; PubMed Central PMCID: PMCPMC7824028.

25. Schulz C, Eschbaumer M, Rudolf M, Konig P, Keller M, Bauer C, et al. Experimental infection of South American camelids with bluetongue virus serotype 8. Vet Microbiol. 2012;154(3-4):257–65. Epub 2011/08/25. doi: 10.1016/j.vetmic.2011.07.025. PubMed PMID: 21862245.

26. Schulz C, Sailleau C, Breard E, Flannery J, Viarouge C, Zientara S, et al. Experimental infection of sheep, goats and cattle with a bluetongue virus serotype 4 field strain from Bulgaria, 2014. Transbound Emerg Dis. 2018;65(2):e243–e50. Epub 2017/11/10. doi: 10.1111/tbed.12746. PubMed PMID: 29119690.

27. Du Toit R. The transmission of blue-tongue and horse-sickness by Culicoides. 1944.

28. Hourrigan JL, Klingsporn AL. Bluetongue: the disease in cattle. Australian veterinary journal. 1975;51(4):170–4.

29. Hutcheon D. The Malarial Catarrhal Fever of Sheep in South Africa. The Veterinary Journal and Annals of Comparative Pathology. 1893;37(11):330–4. doi: https://doi.org/10.1016/S2543-3377(17)49810-4.

30. Owen NC, Du Toit RM, Howell PG. Bluetongue in cattle: typing of viruses isolated from cattle exposed to natural infections. Onderstepoort J Vet Res. 1965;32(1):3–6. Epub 1965/06/01. PubMed PMID: 4960029.

31. Spreull J. Malarial catarrhal fever (Bluetongue) of sheep in South Africa. J Comp Pathol. 1905;18:321–37.

32. Caporale M, Di Gialleonorado L, Janowicz A, Wilkie G, Shaw A, Savini G, et al. Virus and host factors affecting the clinical outcome of bluetongue virus infection. J Virol. 2014;88(18):10399–411. Epub 2014/07/06. doi: 10.1128/JVI.01641-14. PubMed PMID: 24991012; PubMed Central PMCID: PMCPMC4178883.

33. Darpel KE, Batten CA, Veronesi E, Shaw AE, Anthony S, Bachanek-Bankowska K, et al. Clinical signs and pathology shown by British sheep and cattle infected with bluetongue virus serotype 8 derived from the 2006 outbreak in northern Europe. Vet Rec. 2007;161(8):253–61. Epub 2007/08/28. doi: 10.1136/vr.161.8.253. PubMed PMID: 17720961.

34. Erasmus BJ. Bluetongue virus. Virus infections of ruminants. 1990;3:227–37.

35. Verwoerd DW, Erasmus BJ. Bluetongue. In: Coetzer JAW, Tustin RC, editors. Infectious Diseases of Livestock. 2nd ed. Cape Town, South Africa: Oxford University Press; 2004. p. 1201–20.

36. DeMaula CD, Leutenegger CM, Bonneau KR, MacLachlan NJ. The role of endothelial cell-derived inflammatory and vasoactive mediators in the pathogenesis of bluetongue. Virology. 2002;296(2):330–7. Epub 2002/06/19. doi: 10.1006/viro.2002.1476. PubMed PMID: 12069531.

37. DeMaula CD, Leutenegger CM, Jutila MA, MacLachlan NJ. Bluetongue virus-induced activation of primary bovine lung microvascular endothelial cells. Vet Immunol Immunopathol. 2002;86(3-4):147–57. Epub 2002/05/15. doi: 10.1016/s0165-2427(02)00012-0. PubMed PMID: 12007881.

38. Melzi E, Caporale M, Rocchi M, Martin V, Gamino V, di Provvido A, et al. Follicular dendritic cell disruption as a novel mechanism of virus-induced immunosuppression. Proc Natl Acad Sci U S A. 2016;113(41):E6238–E47. Epub 2016/09/28. doi: 10.1073/pnas.1610012113. PubMed PMID: 27671646; PubMed Central PMCID: PMCPMC5068271.

39. Schwartz-Cornil I, Mertens PP, Contreras V, Hemati B, Pascale F, Breard E, et al. Bluetongue virus: virology, pathogenesis and immunity. Vet Res. 2008;39(5):46. Epub 2008/05/23. doi: 10.1051/vetres:2008023. PubMed PMID: 18495078.

40. Maclachlan NJ, Drew CP, Darpel KE, Worwa G. The pathology and pathogenesis of bluetongue. J Comp Pathol. 2009;141(1):1–16. Epub 2009/05/30. doi: 10.1016/j.jcpa.2009.04.003. PubMed PMID: 19476953.

41. Bekker J, De Kock G, Quinlan J. The occurrence and identification of bluetongue in cattle-the so-called pseudo-foot and mouth disease in South Africa. 1934.

42. Dal Pozzo F, De Clercq K, Guyot H, Vandemeulebroucke E, Sarradin P, Vandenbussche F, et al. Experimental reproduction of bluetongue virus serotype 8 clinical disease in calves. Vet Microbiol. 2009;136(3-4):352–8. Epub 2009/01/09. doi: 10.1016/j.vetmic.2008.11.012. PubMed PMID: 19128895.

43. MacLachlan NJ, Fuller FJ. Genetic stability in calves of a single strain of bluetongue virus. Am J Vet Res. 1986;47(4):762–4. Epub 1986/04/01. PubMed PMID: 3008608.

44. MacLachlan NJ, Heidner HW, Fuller FJ. Humoral immune response of calves to bluetongue virus infection. Am J Vet Res. 1987;48(7):1031–5. Epub 1987/07/01. PubMed PMID: 2820272.

45. MacLachlan NJ, Jagels G, Rossitto PV, Moore PF, Heidner HW. The pathogenesis of experimental bluetongue virus infection of calves. Vet Pathol. 1990;27(4):223–9. Epub 1990/07/01. doi: 10.1177/030098589002700402. PubMed PMID: 2169663.

46. Parsonson IM, Della-Porta AJ, McPhee DA, Cybinski DH, Squire KR, Uren MF. Experimental infection of bulls and cows with bluetongue virus serotype 20. Aust Vet J. 1987;64(1):10–3. Epub 1987/01/01. doi: 10.1111/j.1751-0813.1987.tb06048.x. PubMed PMID: 3036056.

47. Parsonson IM, Della-Porta AJ, McPhee DA, Cybinski DH, Squire KR, Uren MF. Bluetongue virus serotype 20: experimental infection of pregnant heifers. Aust Vet J. 1987;64(1):14–7. Epub 1987/01/01. doi: 10.1111/j.1751-0813.1987.tb06049.x. PubMed PMID: 3036057.

48. Foster NM, Luedke AJ, Parsonson IM, Walton TE. Temporal relationships of viremia, interferon activity, and antibody responses of sheep infected with several bluetongue virus strains. Am J Vet Res. 1991;52(2):192–6. Epub 1991/02/01. PubMed PMID: 1707246.

49. Fulton RW, Pearson NJ. Interferon induction in bovine and feline monolayer cultures by four bluetongue virus serotypes. Can J Comp Med. 1982;46(1):100–2. Epub 1982/01/01. PubMed PMID: 6176301; PubMed Central PMCID: PMCPMC1320205.

50. Huismans H. Bluetongue virus-induced interferon synthesis. Onderstepoort J Vet Res. 1969;36(2):181–5. Epub 1969/12/01. PubMed PMID: 4340875.

51. MacLachlan NJ, Schore CE, Osburn BI. Antiviral responses of bluetongue virus-inoculated bovine fetuses and their dams. Am J Vet Res. 1984;45(7):1469–73. Epub 1984/07/01. PubMed PMID: 24049920.

52. MacLachlan NJ, Thompson J. Bluetongue virus-induced interferon in cattle. Am J Vet Res. 1985;46(6):1238–41. Epub 1985/06/01. PubMed PMID: 2411172.

53. Russell H, O’Toole DT, Bardsley K, Davis WC, Ellis JA. Comparative effects of bluetongue virus infection of ovine and bovine endothelial cells. Vet Pathol. 1996;33(3):319–31. Epub 1996/05/01. doi: 10.1177/030098589603300309. PubMed PMID: 8740706.

54. Lin Q, Meloni D, Pan Y, Xia M, Rodgers J, Shepard S, et al. Enantioselective synthesis of Janus kinase inhibitor INCB018424 via an organocatalytic aza-Michael reaction. Org Lett. 2009;11(9):1999–2002. Epub 2009/04/24. doi: 10.1021/ol900350k. PubMed PMID: 19385672.

55. Quintas-Cardama A, Vaddi K, Liu P, Manshouri T, Li J, Scherle PA, et al. Preclinical characterization of the selective JAK1/2 inhibitor INCB018424: therapeutic implications for the treatment of myeloproliferative neoplasms. Blood. 2010;115(15):3109–17. Epub 2010/02/05. doi: 10.1182/blood-2009-04-214957. PubMed PMID: 20130243; PubMed Central PMCID: PMCPMC3953826.

56. Schoggins JW, Wilson SJ, Panis M, Murphy MY, Jones CT, Bieniasz P, et al. A diverse range of gene products are effectors of the type I interferon antiviral response. Nature. 2011;472(7344):481–5. Epub 2011/04/12. doi: 10.1038/nature09907. PubMed PMID: 21478870; PubMed Central PMCID: PMCPMC3409588.

57. Rihn SJ, Aziz MA, Stewart DG, Hughes J, Turnbull ML, Varela M, et al. TRIM69 Inhibits Vesicular Stomatitis Indiana Virus. J Virol. 2019;93(20). Epub 2019/08/04. doi: 10.1128/JVI.00951-19. PubMed PMID: 31375575; PubMed Central PMCID: PMCPMC6798119.

58. Wickenhagen A, Sugrue E, Lytras S, Kuchi S, Noerenberg M, Turnbull ML, et al. A prenylated dsRNA sensor protects against severe COVID-19. Science. 2021;374(6567):eabj3624. Epub 2021/09/29. doi: 10.1126/science.abj3624. PubMed PMID: 34581622.

59. Pinto RM, Bakshi S, Lytras S, Zakaria MK, Swingler S, Worrell JC, et al. Zoonotic avian influenza viruses evade human BTN3A3 restriction. bioRxiv. 2022:2022.06.14.496196. doi: 10.1101/2022.06.14.496196.

60. Shaw AE, Hughes J, Gu Q, Behdenna A, Singer JB, Dennis T, et al. Fundamental properties of the mammalian innate immune system revealed by multispecies comparison of type I interferon responses. PLoS Biol. 2017;15(12):e2004086. Epub 2017/12/19. doi: 10.1371/journal.pbio.2004086. PubMed PMID: 29253856; PubMed Central PMCID: PMCPMC5747502.

61. Kane M, Zang TM, Rihn SJ, Zhang F, Kueck T, Alim M, et al. Identification of Interferon-Stimulated Genes with Antiretroviral Activity. Cell Host Microbe. 2016;20(3):392–405. Epub 2016/09/16. doi: 10.1016/j.chom.2016.08.005. PubMed PMID: 27631702; PubMed Central PMCID: PMCPMC5026698.

62. Kang D, Gao S, Tian Z, Huang D, Guan G, Liu G, et al. Ovine viperin inhibits bluetongue virus replication. Mol Immunol. 2020;126:87–94. Epub 2020/08/14. doi: 10.1016/j.molimm.2020.07.014. PubMed PMID: 32784101.

63. !!! INVALID CITATION !!! [61, 62].

64. Deliu LP, Ghosh A, Grewal SS. Investigation of protein synthesis in Drosophila larvae using puromycin labelling. Biol Open. 2017;6(8):1229–34. Epub 2017/06/24. doi: 10.1242/bio.026294. PubMed PMID: 28642244; PubMed Central PMCID: PMCPMC5576084.

65. Gillespie M, Jassal B, Stephan R, Milacic M, Rothfels K, Senff-Ribeiro A, et al. The reactome pathway knowledgebase 2022. Nucleic Acids Res. 2022;50(D1):D687–D92. Epub 2021/11/18. doi: 10.1093/nar/gkab1028. PubMed PMID: 34788843; PubMed Central PMCID: PMCPMC8689983.

66. Sidiropoulos K, Viteri G, Sevilla C, Jupe S, Webber M, Orlic-Milacic M, et al. Reactome enhanced pathway visualization. Bioinformatics. 2017;33(21):3461–7. Epub 2017/10/28. doi: 10.1093/bioinformatics/btx441. PubMed PMID: 29077811; PubMed Central PMCID: PMCPMC5860170.

67. Takaoka A, Yanai H. Interferon signalling network in innate defence. Cell Microbiol. 2006;8(6):907–22. Epub 2006/05/10. doi: 10.1111/j.1462-5822.2006.00716.x. PubMed PMID: 16681834.

68. Roberts RM, Liu L, Guo Q, Leaman D, Bixby J. The evolution of the type I interferons. J Interferon Cytokine Res. 1998;18(10):805–16. Epub 1998/11/11. doi: 10.1089/jir.1998.18.805. PubMed PMID: 9809615.

69. Peters SO, Hussain T, Adenaike AS, Hazzard J, Morenikeji OB, De Donato M, et al. Evolutionary Pattern of Interferon Alpha Genes in Bovidae and Genetic Diversity of IFNAA in the Bovine Genome. Front Immunol. 2020;11:580412. Epub 2020/10/30. doi: 10.3389/fimmu.2020.580412. PubMed PMID: 33117386; PubMed Central PMCID: PMCPMC7561390.

70. McCarthy DJ, Chen Y, Smyth GK. Differential expression analysis of multifactor RNA-Seq experiments with respect to biological variation. Nucleic Acids Res. 2012;40(10):4288–97. Epub 2012/01/31. doi: 10.1093/nar/gks042. PubMed PMID: 22287627; PubMed Central PMCID: PMCPMC3378882.

71. Robinson MD, McCarthy DJ, Smyth GK. edgeR: a Bioconductor package for differential expression analysis of digital gene expression data. Bioinformatics. 2010;26(1):139–40. Epub 2009/11/17. doi: 10.1093/bioinformatics/btp616. PubMed PMID: 19910308; PubMed Central PMCID: PMCPMC2796818.

72. Iwasaki A. A virological view of innate immune recognition. Annu Rev Microbiol. 2012;66:177–96. Epub 2012/09/22. doi: 10.1146/annurev-micro-092611-150203. PubMed PMID: 22994491; PubMed Central PMCID: PMCPMC3549330.

73. Diamond MS, Farzan M. The broad-spectrum antiviral functions of IFIT and IFITM proteins. Nature reviews Immunology. 2013;13(1):46–57. Epub 2012/12/15. doi: 10.1038/nri3344. PubMed PMID: 23237964; PubMed Central PMCID: PMCPMC3773942.

74. Rivera-Serrano EE, Gizzi AS, Arnold JJ, Grove TL, Almo SC, Cameron CE. Viperin Reveals Its True Function. Annu Rev Virol. 2020;7(1):421–46. Epub 2020/07/01. doi: 10.1146/annurev-virology-011720-095930. PubMed PMID: 32603630; PubMed Central PMCID: PMCPMC8191541.

75. Hoang HD, Graber TE, Alain T. Battling for Ribosomes: Translational Control at the Forefront of the Antiviral Response. J Mol Biol. 2018;430(14):1965–92. Epub 2018/05/11. doi: 10.1016/j.jmb.2018.04.040. PubMed PMID: 29746850.

76. Stern-Ginossar N, Thompson SR, Mathews MB, Mohr I. Translational Control in Virus-Infected Cells. Cold Spring Harb Perspect Biol. 2019;11(3). Epub 2018/06/13. doi: 10.1101/cshperspect.a033001. PubMed PMID: 29891561; PubMed Central PMCID: PMCPMC6396331.

77. Walsh D, Mohr I. Viral subversion of the host protein synthesis machinery. Nat Rev Microbiol. 2011;9(12):860–75. Epub 2011/10/18. doi: 10.1038/nrmicro2655. PubMed PMID: 22002165; PubMed Central PMCID: PMCPMC7097311.

78. Boyce M, Celma CC, Roy P. Bluetongue virus non-structural protein 1 is a positive regulator of viral protein synthesis. Virol J. 2012;9:178. Epub 2012/08/31. doi: 10.1186/1743-422X-9-178. PubMed PMID: 22931514; PubMed Central PMCID: PMCPMC3479040.

79. Chauveau E, Doceul V, Lara E, Breard E, Sailleau C, Vidalain PO, et al. NS3 of bluetongue virus interferes with the induction of type I interferon. J Virol. 2013;87(14):8241–6. Epub 2013/05/10. doi: 10.1128/JVI.00678-13. PubMed PMID: 23658442; PubMed Central PMCID: PMCPMC3700197.

80. Ratinier M, Caporale M, Golder M, Franzoni G, Allan K, Nunes SF, et al. Identification and characterization of a novel non-structural protein of bluetongue virus. PLoS Pathog. 2011;7(12):e1002477. Epub 2012/01/14. doi: 10.1371/journal.ppat.1002477. PubMed PMID: 22241985; PubMed Central PMCID: PMCPMC3248566.

81. Ratinier M, Shaw AE, Barry G, Gu Q, Di Gialleonardo L, Janowicz A, et al. Bluetongue Virus NS4 Protein Is an Interferon Antagonist and a Determinant of Virus Virulence. J Virol. 2016;90(11):5427–39. doi: 10.1128/JVI.00422-16. PubMed PMID: 27009961; PubMed Central PMCID: PMC4934764.

82. Li Z, Lu D, Yang H, Li Z, Zhu P, Xie J, et al. Bluetongue virus non-structural protein 3 (NS3) and NS4 coordinatively antagonize type I interferon signaling by targeting STAT1. Vet Microbiol. 2021;254:108986. Epub 2021/01/25. doi: 10.1016/j.vetmic.2021.108986. PubMed PMID: 33486325.

83. Arnaud F, Black SG, Murphy L, Griffiths DJ, Neil SJ, Spencer TE, et al. Interplay between ovine bone marrow stromal cell antigen 2/tetherin and endogenous retroviruses. J Virol. 2010;84(9):4415–25. Epub 2010/02/26. doi: 10.1128/JVI.00029-10. PubMed PMID: 20181686; PubMed Central PMCID: PMCPMC2863748.

84. Varela M, Schnettler E, Caporale M, Murgia C, Barry G, McFarlane M, et al. Schmallenberg virus pathogenesis, tropism and interaction with the innate immune system of the host. PLoS Pathog. 2013;9(1):e1003133. Epub 2013/01/18. doi: 10.1371/journal.ppat.1003133. PubMed PMID: 23326235; PubMed Central PMCID: PMCPMC3542112.

85. Smith MC, Goddard ET, Perusina Lanfranca M, Davido DJ. hTERT extends the life of human fibroblasts without compromising type I interferon signaling. PLoS One. 2013;8(3):e58233. Epub 2013/03/09. doi: 10.1371/journal.pone.0058233. PubMed PMID: 23472163; PubMed Central PMCID: PMCPMC3589264.

86. Charman M, McFarlane S, Wojtus JK, Sloan E, Dewar R, Leeming G, et al. Constitutive TRIM22 Expression in the Respiratory Tract Confers a Pre-Existing Defence Against Influenza A Virus Infection. Front Cell Infect Microbiol. 2021;11:689707. Epub 2021/10/09. doi: 10.3389/fcimb.2021.689707. PubMed PMID: 34621686; PubMed Central PMCID: PMCPMC8490869.

87. Boyce M, Celma CC, Roy P. Development of reverse genetics systems for bluetongue virus: recovery of infectious virus from synthetic RNA transcripts. J Virol. 2008;82(17):8339–48. Epub 2008/06/20. doi: 10.1128/JVI.00808-08. PubMed PMID: 18562540; PubMed Central PMCID: PMCPMC2519640.

88. Boyce M, Roy P. Recovery of infectious bluetongue virus from RNA. J Virol. 2007;81(5):2179–86. Epub 2006/12/08. doi: 10.1128/JVI.01819-06. PubMed PMID: 17151117; PubMed Central PMCID: PMCPMC1865916.

89. Reed LJ, Muench H. A simple method of estimating fifty per cent endpoints. American journal of epidemiology. 1938;27(3):493–7.

90. Spearman C. The method of right and wrong cases (constant stimuli) without Gauss’s formulae. British journal of psychology. 1908;2(3):227–42.

91. Bakshi S, Taylor J, Strickson S, McCartney T, Cohen P. Identification of TBK1 complexes required for the phosphorylation of IRF3 and the production of interferon beta. Biochem J. 2017;474(7):1163–74. Epub 2017/02/06. doi: 10.1042/BCJ20160992. PubMed PMID: 28159912; PubMed Central PMCID: PMCPMC5350611.

92. Labun K, Montague TG, Krause M, Torres Cleuren YN, Tjeldnes H, Valen E. CHOPCHOP v3: expanding the CRISPR web toolbox beyond genome editing. Nucleic Acids Res. 2019;47(W1):W171–W4. Epub 2019/05/21. doi: 10.1093/nar/gkz365. PubMed PMID: 31106371; PubMed Central PMCID: PMCPMC6602426.

93. Sanjana NE, Shalem O, Zhang F. Improved vectors and genome-wide libraries for CRISPR screening. Nat Methods. 2014;11(8):783–4. Epub 2014/07/31. doi: 10.1038/nmeth.3047. PubMed PMID: 25075903; PubMed Central PMCID: PMCPMC4486245.

94. Hofmann M, Griot C, Chaignat V, Perler L, Thur B. [Bluetongue disease reaches Switzerland]. Schweiz Arch Tierheilkd. 2008;150(2):49–56. Epub 2008/03/29. doi: 10.1024/0036-7281.150.2.49. PubMed PMID: 18369049.

95. Liao Y, Smyth GK, Shi W. featureCounts: an efficient general purpose program for assigning sequence reads to genomic features. Bioinformatics. 2014;30(7):923–30. Epub 2013/11/15. doi: 10.1093/bioinformatics/btt656. PubMed PMID: 24227677.

96. Yu G, He Q-Y. ReactomePA: an R/Bioconductor package for reactome pathway analysis and visualization. Molecular BioSystems. 2016;12(2):477–9. doi: 10.1039/C5MB00663E.

97. Durinck S, Spellman PT, Birney E, Huber W. Mapping identifiers for the integration of genomic datasets with the R/Bioconductor package biomaRt. Nat Protoc. 2009;4(8):1184–91. Epub 2009/07/21. doi: 10.1038/nprot.2009.97. PubMed PMID: 19617889; PubMed Central PMCID: PMCPMC3159387.

